# Degeneracy explains diversity in interneuronal regulation of pattern separation in heterogeneous dentate gyrus networks

**DOI:** 10.1101/2025.02.14.638282

**Authors:** Sarang Saini, Rishikesh Narayanan

## Abstract

**Background and motivations:** The Marr-Albus theory postulates that pattern separation is realized by divergent feedforward excitatory connectivity. Yet, there are several lines of evidence for strong but differential regulation of pattern separation by local circuit connectivity, even when feedforward connectivity is divergent. What are the relative contributions of divergent feedforward connectivity and local circuit interactions to pattern separation? How do we reconcile the contrasting lines of evidence on local circuit regulation of pattern separation in circuits endowed with divergent feedforward connectivity? In this study, we quantitatively address these questions in a population of heterogeneous dentate gyrus (DG) networks, where we enforced feedforward connectivity to be identically divergent.

**Methodology:** We generated tens of thousands of random spiking neuronal models to arrive at thousands of non-repeating heterogeneous single-neuron models of four different DG neuronal subtypes, each satisfying their respective functional characteristics. We connected these heterogeneous populations of neurons with subtype proportions and local connectivity that reflected the DG microcircuit. In a second level of unbiased search, we generated 20,000 identical networks that differed from each other only in their synaptic weight values. Thus, within the Marr-Albus framework, these networks that were identical in terms of neuronal composition and divergent feedforward connectivity should all be capable of performing effective pattern separation. To test this, we fed these networks with morphed sets of input patterns, recorded granule cell outputs, and computed similarity metrics based on correlation measures across input or output patterns. We developed novel quantitative metrics for pattern separation from plots of output similarity *vs.* input similarity and validated each network with bounds on these metrics.

**Results:** Despite being identical in terms of divergent feedforward connectivity, we found only a very small proportion (47 of 20,000 or 0.23%) of the randomly generated networks to manifest effective pattern separation. We tested the specific contributions of interneurons by assessing pattern separation in all pattern-separating networks after individually deleting each of the three interneuron subtypes. Strikingly, we found pronounced network-to-network variability in how each interneuron subtype contributed to granule cell sparsity and pattern separation. We traced this variability to differences in local synaptic connectivity, which also resulted in network-to-network variability in firing rates and sparsity of different interneurons. Finally, we found heterogeneous DG networks to be more resilient to synaptic jitter compared to their homogeneous counterparts, with specific reference to pattern separation computed through average firing rate correlations.

**Implications:** Our population-of-networks approach clearly shows that divergent connectivity of afferent inputs does not guarantee pattern separation in DG networks. Instead, we demonstrate strong yet variable roles for interneurons and local connectivity in implementing pattern separation. Importantly, our analyses unveil degeneracy in DG circuits, whereby similar pattern separation efficacy was achieved through disparate local-circuit interactions. These observations, alongside network-to-network variability in dependencies on different interneurons, strongly advocate the complex adaptive systems approach as a unifying framework to study DG pattern separation.

## INTRODUCTION

Pattern separation is the ability of a system to discriminate between similar inputs by transforming them into distinct outputs. In the context of the brain, inputs could originate from similar behavioral contexts or experiences and pattern separation is accomplished by neural circuits that transform these similar inputs into dissimilar neuronal outputs. The dentate gyrus (DG), the gateway to the hippocampus proper, is a well-studied pattern separation circuit that has also been implicated in memory formation, spatial navigation, and different learning paradigms. Historically, DG was postulated as a pattern-separating network based on divergent excitatory afferent connectivity from the entorhinal cortex (Albus, 1971; Marr, 1971). Since then, several studies have provided strong lines of evidence for a role of DG in pattern separation (Leutgeb et al., 2007; McHugh et al., 2007; Bakker et al., 2008; Aimone et al., 2014; Knierim and Neunuebel, 2016; Anacker and Hen, 2017; Scharfman, 2025). Despite this, the mechanisms underlying pattern separation in the dentate gyrus have been highly debated (Cayco-Gajic and Silver, 2019; Borzello et al., 2023), especially regarding the origins of sparse activity of granule cells (Dieni et al., 2013; Diamantaki et al., 2016; Danielson et al., 2017; GoodSmith et al., 2017; Senzai and Buzsaki, 2017; Severa et al., 2017), the roles of the several DG local-circuit components (Mott et al., 1997; Sik et al., 1997; Bartos et al., 2001; Amaral et al., 2007; Houser, 2007; Stefanelli et al., 2016; Hainmueller and Bartos, 2018; Elgueta and Bartos, 2019; Hainmueller et al., 2024; Scharfman, 2025), implications for the widespread heterogeneities in the DG circuit (Mishra and Narayanan, 2019; Mishra and Narayanan, 2020; Huckleberry and Shansky, 2021), and the impact of adult neurogenesis and hyperplasticity on circuit formation (Schmidt-Hieber et al., 2004; Ge et al., 2007; Aimone et al., 2014; Abrous and Wojtowicz, 2015; Li et al., 2017; Lodge and Bischofberger, 2019; Denoth-Lippuner and Jessberger, 2021; Shridhar et al., 2022). There are conflicting reports especially on the specific roles of the different interneurons in the intricately connected DG circuitry in pattern separation, sparsity, and other functions (Dieni et al., 2013; Stefanelli et al., 2016; Chavlis et al., 2017; Hainmueller and Bartos, 2018; Sun et al., 2020; Guzman et al., 2021; Yen et al., 2022; Borzello et al., 2023; Hainmueller et al., 2024).

What are the relative contributions of divergent feedforward connectivity (as postulated by the Marr-Albus theory) and local circuit interactions (involving conflicting reports) to pattern separation? How do we reconcile the contrasting lines of evidence on interneuron regulation of pattern separation in circuits endowed with divergent feedforward connectivity? Is there a unifying framework that could synthesize these conflicting lines of evidence on the roles of different circuit components in executing pattern separation? In addressing these questions, we chose an exhaustive population-of-networks approach to study pattern separation in the heterogeneous DG circuit that was endowed with different neuronal subtypes. First, we generated tens of thousands of random spiking neuronal models to arrive at thousands of non-repeating single-neuron models of four different DG neuronal subtypes, each satisfying their respective functional characteristics. We connected these heterogeneous populations of neurons with subtype proportions and local connectivity that reflected the DG microcircuit. In a second level of unbiased search, we generated 20,000 identical networks that differed from each other only in their synaptic weight values and found only 47 of these networks to perform pattern separation.

We found the synaptic weight values in these heterogeneous pattern-separating networks to span a large range with weak pairwise relationships between weight values across networks. These observations unveiled the manifestation of degeneracy in these networks, whereby pattern separation emerged through interactions among several non-unique, non-random combinations of DG circuit elements. Strikingly, our analyses unveiled pronounced network-to-network variability in the impact of deleting each interneuron subtype on pattern separation performance, sparsity, and firing rates. We attributed these differences to the large range of local-circuit synaptic weight values across different networks that achieved pattern separation through degenerate circuit connectivity. Finally, our analyses also demonstrated that DG networks endowed with within-cell-type heterogeneities were more resilient to synaptic perturbations compared to their homogeneous counterparts.

Together, our population-of-networks approach clearly shows that divergent connectivity of afferent inputs does not guarantee pattern separation in DG networks. Instead, we demonstrate strong yet variable roles for interneurons and local connectivity in implementing pattern separation. Importantly, our analyses unveil multi-scale degeneracy in DG circuits, whereby similar single-neuron function was achieved through disparate parametric combinations and similar pattern separation efficacy was implemented by disparate circuit interactions. Our demonstration that pattern separation emerged through interactions among several non-unique, non-random combinations of DG circuit elements strongly advocate the use of the complex systems framework to study DG pattern separation.

## RESULTS

### Functionally validated heterogeneous populations of models for each of the four different neuronal subtypes

We generated four independent heterogeneous populations of granule cells (GC), basket cells (BC), mossy cells (MC), and HIPP cells (HC). We employed the well-established multi-parametric multi-objective stochastic search (MPMOSS) algorithm (Foster et al., 1993; Prinz et al., 2003; Taylor et al., 2009; Marder and Taylor, 2011; Rathour and Narayanan, 2012; Rathour and Narayanan, 2014; Mishra and Narayanan, 2019; Mishra and Narayanan, 2021c; Shridhar et al., 2022; Sinha and Narayanan, 2022; Schneider et al., 2023; Kumari and Narayanan, 2024) to generate valid heterogeneous populations of each cell type through independent and unbiased stochastic searches (Fig. 1*A*). We used adaptive exponential integrate-and-fire (aEIF) spiking neuron models for all four neuronal subtypes. The search spanned the same set of 9 parameters, but with different ranges, for each of the different cell types (Supplementary Table S1–S4). Randomly generated models were validated against subtype-specific signature electrophysiological measurements (Supplementary Table S5). We used six subtype-specific electrophysiological measurements for validation (Lübke et al., 1998; Bartos et al., 2001; Ratzliff et al., 2002; Krueppel et al., 2011; Mishra and Narayanan, 2019): membrane time constant (*τ_m_*), sag ratio (*sag*), input resistance (*R_in_*), spike frequency adaptation (*SFA*), and action potential firing frequency for 50 pA (*f*_50_) or 150 pA (*f*_150_) pulse current injections. We randomly generated 100,000 granule cells, 60,000 basket cells, 10,000 mossy cells, and 20,000 HIPP cells from respective parametric distributions and validated them against their signature electrophysiological characteristics. Of these models, we found 6378 granule cells (∼6.3% of the total generated models), 573 basket cells (∼0.9%), 273 mossy cells (∼2.7%), and 87 HIPP cells (∼0.4%) to satisfy all electrophysiological constraints (Fig. 1*B*). We declared these as valid models belonging to the specific neuronal subtypes.

**Figure 1.**
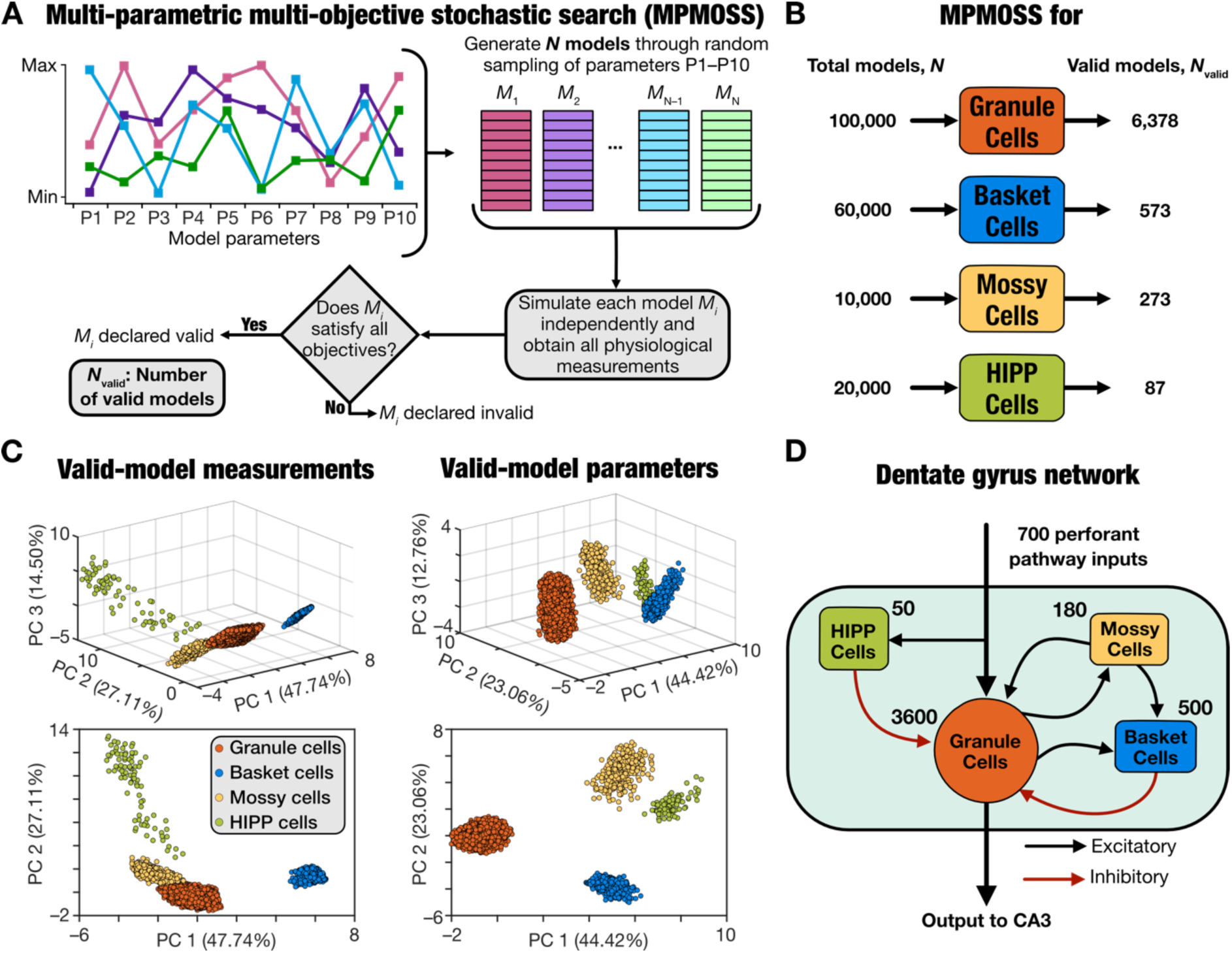
Dentate gyrus networks were constructed with cell-type-specific heterogeneous cell populations derived from independent unbiased stochastic searches. (A) Schematic of the MPMOSS algorithm used for generating heterogeneous populations of the four different cell types that were used to construct the heterogeneous dentate gyrus (DG) network. To ensure that cellular heterogeneity isn’t constricted by searches spanning confined subspaces, we used unbiased stochastic searches spanning a wide parametric space, independently performed for each cell type. For each cell type, the 9-dimensional parametric spaces of the aEIF models were randomly sampled (Supplementary Table S1–S4) to yield *N* individual models. Each randomly generated model was then validated against the physiological properties of the respective cell type (Supplementary Table S5). Models that satisfied all physiological properties for that cell types were declared to be valid, with *N_valid_* representing the number of such valid models. (B) The total number of random models that were generated (*N*) and the number of valid models (*N_valid_*) obtained for each of the four cell types are shown. The number of valid models generated was constrained by the number of distinct models of each cell type required for generating the heterogeneous DG network. (C) All cell types were modelled as aEIF models with searches spanning the same 10 model parameters and were validated against the same set of 6 physiological measurements. Therefore, visualization of all valid models of the four different cell types (from panel B) on the same reduced 3D space was feasible through principal component analysis spanning all valid models. The clustering across and the heterogeneities within different cell types may be noted in both the measurement (*left*) as well as parametric (*right*) spaces. The numbers along each principal component (PC) axis indicates the percentage variance explained by that dimension. PC1 *vs*. PC2 is plotted below to demonstrate distinct clusters. (D) A graphical representation of the heterogeneous DG network architecture with 3600 unique granule cells and three unique sets of interneuron subtypes (500 basket, 180 mossy, and 50 HIPP cells) randomly picked from the respective heterogeneous population. The network receives inputs from the perforate pathway and send its outputs to CA3 pyramidal cells.

We confirmed heterogeneities in the model parameters (Supplementary Fig. S1) and physiological measurements (Supplementary Fig. S2) of these models by noting the widespread nature of the ranges of each parameter and measurement, spanning their respective valid ranges in entirety. In addition, we plotted pairwise dependencies across model parameters (Supplementary Fig. S1) and measurements (Supplementary Fig. S2) and found most pairwise correlations to be weak. These observations are also consistent with other computational and electrophysiological studies demonstrating degeneracy in different DG cell types (Mishra and Narayanan, 2019; Mishra and Narayanan, 2021c, b; Shridhar et al., 2022; Schneider et al., 2023; Kumari and Narayanan, 2024), whereby disparate combinations of model parameters yield signature physiological outcomes. Importantly, since all four neuronal subtypes shared the same parametric and the measurements spaces, we performed principal component analysis on both spaces spanning all neuronal subtypes to visualize these models in reduced dimensions. We visualized all models as coefficients on the first three principal components, which explained ∼90% of the variance (Fig. 1*C*). We also found that the models belonging to the four neuronal subtypes formed distinct clusters while also manifesting heterogeneities within individual clusters (Fig. 1*C*).

Together, these MPMOSS algorithms yielded heterogeneous populations of models that functionally matched each of the four neuronal subtypes. We used these single-neuron model populations to build a heterogeneous DG network (Fig. 1*D*) that was made of non-repeating neurons with connectivity and synaptic properties (Supplementary Tables S6–S7) adopted from their biological counterparts (Geiger et al., 1997; Bartos et al., 2001; Larimer and Strowbridge, 2008; Myers and Scharfman, 2009; Chavlis et al., 2017). The network model thus was built with models that matched the physiological heterogeneities of the different neuronal subtypes with local and afferent connectivity matching anatomical observations (Fig. 1*D*).

### Progressively morphed patterns as network inputs

The perforant pathway served as inputs to the DG network, with 700 afferent inputs projecting onto granule and HIPP cells within the network (Fig. 1*D*). Each of these 700 PP inputs was modeled as an independent spike train, collectively forming one input pattern. The spike trains were randomly generated, with the spike frequency drawn from a Poisson distribution with *λ* = 8 Hz (Fig. 2*A*) and inter-spike intervals drawn from an exponential distribution with *λ* = 125 ms (Fig. 2*B*). A total of 700 spike trains that were constrained by these distributions (10 examples are shown in Fig. 2*C*) were generated to yield one pattern (designated as *P*_0_ in Fig. 2*D*). A similar randomized generation process with a different seed yielded a second independent pattern (designated as *P*_1_ in Fig. 2*D*). These two patterns *P*_0_ and *P*_1_ served as extremes for generating 9 other morphed intermediate patterns that transitioned progressively from *P*_0_towards *P*_1_(Fig. 2*D*). For a pattern *P_β_*, *β* fraction of the total 700 spike trains were randomly selected from *P*_1_and the remaining 1 − *β* fraction from *P*_0_. We varied β progressively from 0.1 to 0.9 in steps of 0.1, resulting in a total of 11 input patterns (*P*_0_, *P*_0.1_, *P*_0.2_, …, *P*_0.9_and *P*_1_).

**Figure 2:**
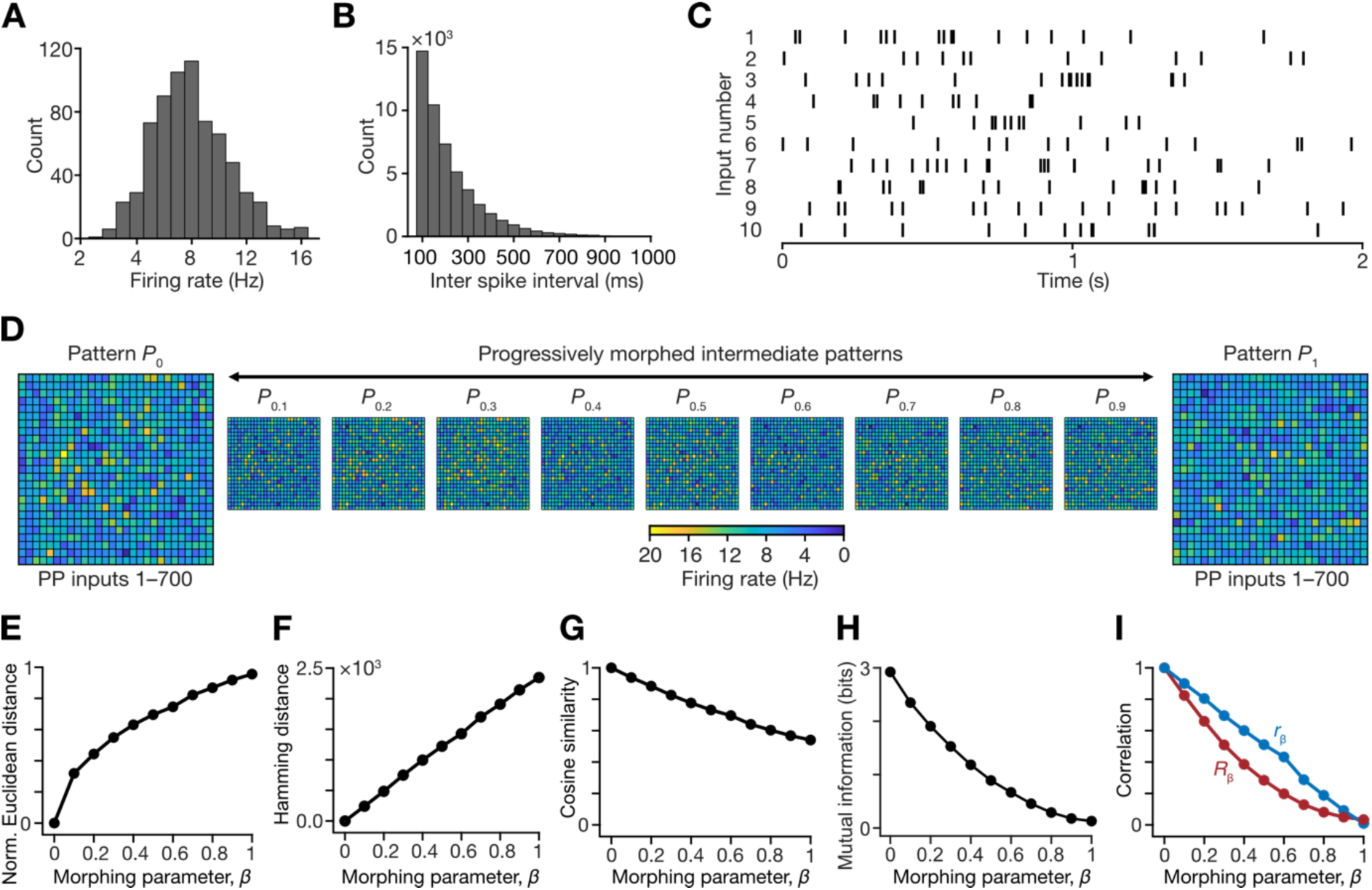
Input structure to the DG network and performance of different distance metrics. (A) Distribution of firing rates of 700 input neurons, derived from a Poisson distribution with a mean of 8 Hz. (B) Distribution of the inter-spike intervals generated from an exponential distribution with *λ* of 125 ms. (C) Raster plot of randomly selected 10 out of 700 perforant pathway input neurons. (D) Two independent firing patterns, *P*_0_and *P*_1_were generated by randomly sampling the firing rate distribution. Each panel shows the heat map representing the average firing rate of 700 (25 × 27) neurons for both the patterns. *P*_0_ was progressively morphed to *P*_1_ to generate 9 intermediate patterns, *P_β_* with 0 < *β* < 1. (E–I) Assessment of the different distance metrics for comparing the 11 different input patterns, with reference to the *P*_0_ pattern. Shown are the computed distances between *P_β_*(with 0 ≤ *β* ≤ 1) and *P*_0_ for normalized Euclidean distance (E), Hamming distance (F), cosine similarity (G), mutual information (H), and the two pattern correlation metrics, *r_β_* and *R_β_* (I).

We assessed the similarity between *P*_0_ and each of the 10 other input patterns using six different metrics: normalized Euclidean distance (Fig. 2*E*), Hamming distance (Fig. 2*F*), cosine similarity (Fig. 2*G*), mutual information (Fig. 2*H*), correlation between average firing rates, *r_β_* (Fig. 2*I*), and the correlation between the instantaneous firing rates (*R_β_*) (Fig. 2*I*). As expected, the two distance-based similarity measurements increased as patterns diverged from each other (Fig. 2*E–F*), while cosine similarity, mutual information, and the correlation-based measures decreased with increasing dissimilarity (Fig. 2*G–I*). The large distances between *P*_0_and *P*_1_, coupled with the gradual changes in the measurement values as functions of the morphing parameter (Fig. 2*E–I*) together confirmed that these patterns could be used to systematically assess pattern separation in our DG network using the morphed inputs approach (Leutgeb et al., 2007; Renno-Costa et al., 2010).

### Correlation-based metrics were invariant to population size, average firing rate, and the form of the firing rate distribution of input patterns

If *S_in_*denoted similarity among the input patterns and *S_out_* represented the same similarity measure computed for the pattern-specific outputs of the network to these inputs, pattern separation was defined when *S_out_* < *S_in_* across different levels of similarity among input patterns (Guzowski et al., 2004; Leutgeb et al., 2007; Renno-Costa et al., 2010; Sahay et al., 2011; Yassa and Stark, 2011; Knierim and Neunuebel, 2016; Borzello et al., 2023). We set three specific criteria in choosing the best of the several similarity measures across patterns made of spike trains (Fig. 3). First, consistent with the divergent anatomical connectivity in the DG network, our model consisted of 700 PP input spike trains and 3600 output spike trains from granule cells (Fig. 1*D*). This difference in number of elements in the input *vs*. the output populations and the need to compare output *vs*. input similarity to quantitatively analyze pattern separation necessitated a metric that was *invariant to population size*. Second, as average firing rates of the input *vs.* output patterns could be very different, we required a metric that was *invariant to the average firing rates* of the population. Third, the firing rate distributions of the input and the output need not obey specific parametrized forms, especially given the sparse nature of GC firing (Jung and McNaughton, 1993; Rolls, 2010, 2013; Diamantaki et al., 2016; Danielson et al., 2017; GoodSmith et al., 2017; Senzai and Buzsaki, 2017; Zhang and Jonas, 2020). Therefore, we laid an additional constraint that the similarity measure should be *invariant to the form of the firing rate distribution*. These three constraints on metrics were essential to ensure that the metric was sensitive solely to the degree of similarity between the two spike trains, thereby enabling effective comparisons between input and output similarity.

**Figure 3:**
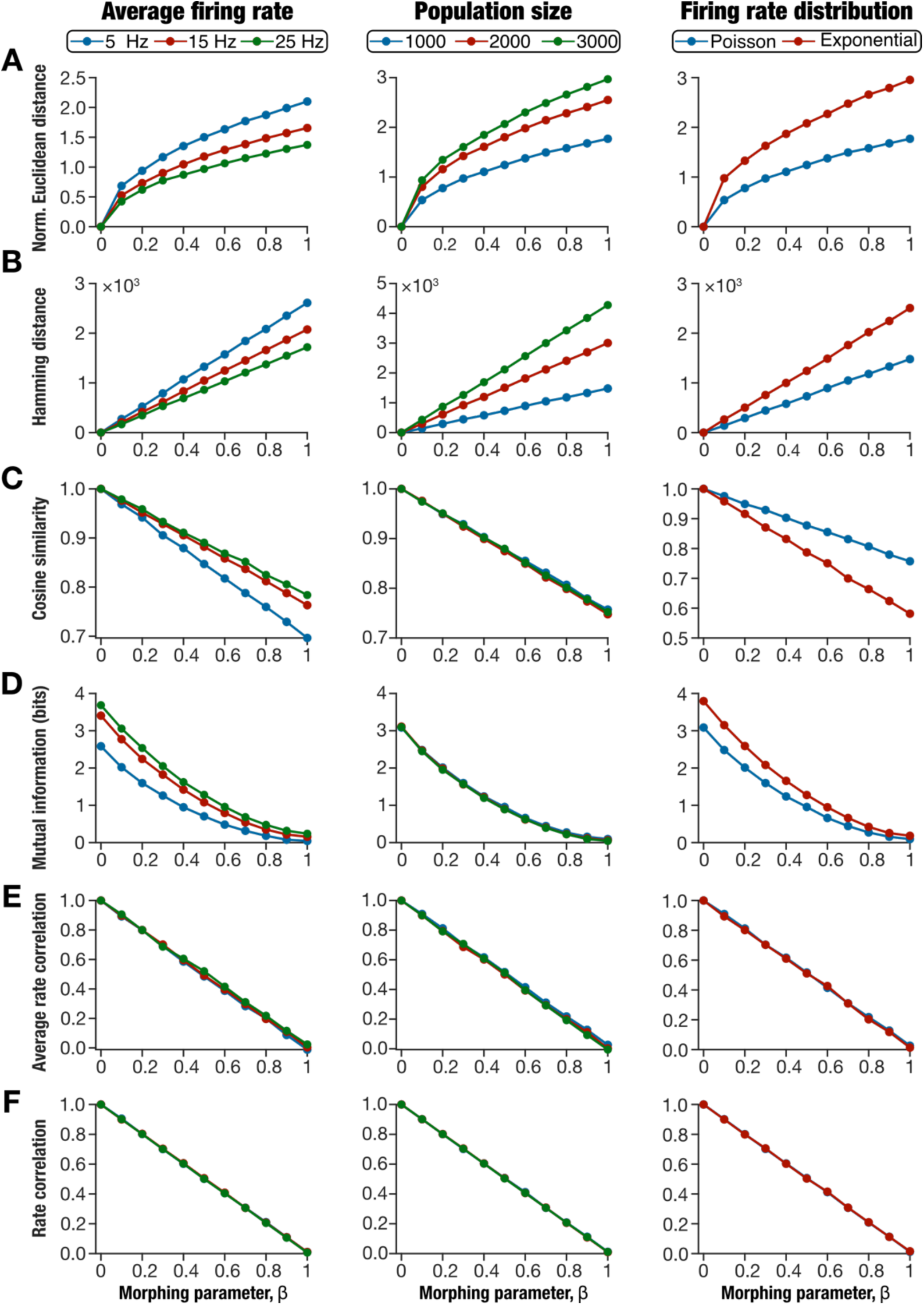
Performance of different metrics used to measure similarity of morphed sets of spike trains. Performance of 6 different metrics that quantified similarity/dissimilarity of spike raster patterns generated for different average firing rate (*first column*), different population sizes (*second column*), and different firing rate distributions (*third column*). In each scenario, ten sets of two distinct patterns (*P*_0_ and *P*_1_) were generated to match the specifications (of firing rate, population size, and distribution type). For each of the 10 *P*_0_–*P*_1_ pairs, nine additional intermediate morphed patterns were created by morphing through *P*_0_ to *P*_1_. The metrics were computed for all patterns with reference to *P*_0_ and the mean values across the ten sets of patterns were plotted as functions of the morphing parameter β. The metrics used were normalized Euclidean distance (A), Hamming distance (B), cosine similarity (C), mutual information (D), average rate correlation *r_β_* (E), and rate correlation *R_β_* (F).

To evaluate the robustness of the different similarity metrics to average firing rates of input patterns, we generated different sets of 11 morphed patterns (as illustrated in Fig. 2), each with a different average firing rate value (5 Hz, 15 Hz, or 25 Hz). To assess invariance to population size, we generated different sets of 11 morphed patterns with 1000, 2000 or 3000 spike trains in each pattern. To examine the robustness of the similarity metrics to different firing rate distributions, we generated two sets of input patterns, one from a Poisson distribution and another from an exponential distribution, with average firing rate set at 8 Hz. For each of these three sets of input patterns, we computed the similarity between *P_β_* and *P*_0_ for each of the six metrics (shown in Fig. 2*E–I*). We plotted the similarity measure values as functions of the morphing parameter *β* for each of the three sets of patterns (Fig. 3).

Across all sets of patterns, all similarity metrics showed monotonic and gradual changes with reference to change in the morphing parameter. We found that Hamming distance, normalized Euclidean distance, cosine similarity, and mutual information measures were sensitive to changes in average firing rate and in firing rate distributions (Fig. 3*A–D*). In addition, the two distance-based metrics were sensitive to changes in population sizes as well (Fig. 3*A–B*). The correlation-based measures, *r_β_* and *R_β_*, were invariant to all the three different tested criteria (Fig. 3*E–F*). Based on these analyses, we chose *r_β_* and *R_β_* as similarity measures for comparing input and output patterns of our network.

### Sparse firing and pattern separation in the default DG network were strongly dependent on interneuron activity

We constructed a heterogeneous DG network by randomly picking non-repeating units of each neuron subtype from their respective valid model populations (Fig. 1), with connectivity and synaptic parameters defined by the biological equivalents (Supplementary Tables S6–S7). We hand-tuned the synaptic weight parameters in this network such that *S_out_* was less than *S_in_*across input patterns, with similarity computed with either the *r_β_* or the *R_β_* metric. This yielded a heterogeneous DG network that executed pattern separation (Fig. 4). In arriving at this network, we presented the 11 morphed inputs patterns to the network and computed the voltage responses of all neurons in the network to arrive at their spike patterns (Fig. 4*A*). We computed the firing rates of all neurons in this default network and found neuron-to-neuron variability across all neuronal subtypes (Fig. 4*B*).

**Figure 4:**
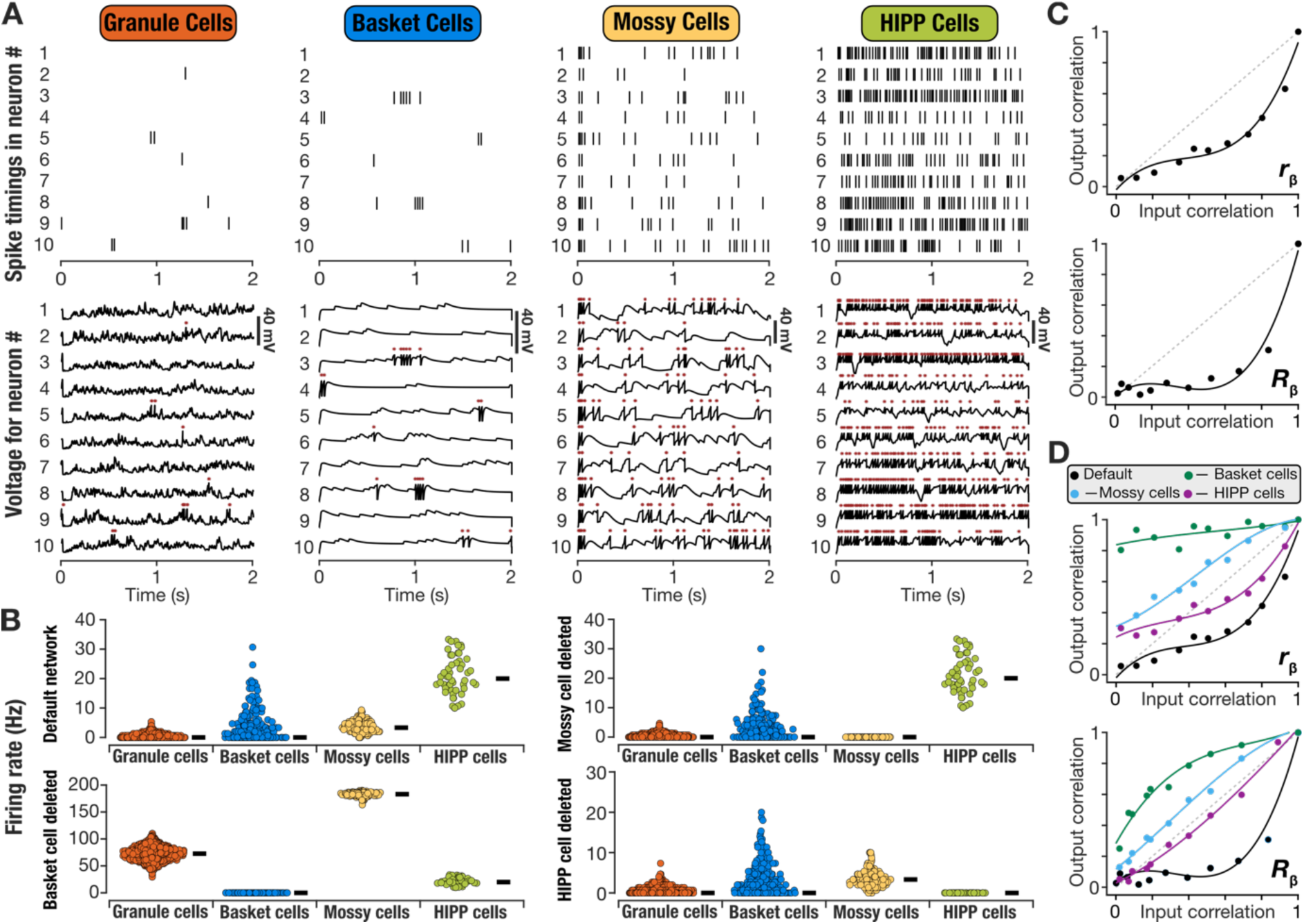
Deletion of different interneuron populations yielded differential impact on pattern separation performance of the default dentate gyrus network model. (A) Raster (*top*) and voltage traces (*bottom*) of 10 different granule, basket, mossy, and HIPP cells of the default DG network presented with the *P*_0_ pattern. Asterisks above voltage traces indicate action potentials, which are represented by the raster plots above. (B) Distribution of firing rates for each neuronal subtype in the default network and in networks where each of the different interneuron subtypes (basket cells, mossy cells, and HIPP cells) were deleted. (C) The ability of the default network to perform pattern separation represented using the plots of output correlation *vs*. input correlation, when correlation was measured using the *r_β_* (*top*) or *R_β_* (*bottom*) metrics. It may be noted that output correlation was lesser than the input correlation, thus signifying the ability of the network to perform effective pattern separation. (D) Pattern separation in the default network and in networks where each of the different interneuron subtypes (basket cells, mossy cells, and HIPP cells) were deleted. Correlation was measured using either the *r_β_* (*top*) or *R_β_* (*bottom*) metrics. For panels C–D, the dots represent the input-output pairs when the network was presented with the 11 morphed patterns and the traces represent a cubic polynomial fitted to these data points.

We found granule cells to show low firing rates and sparse firing, with many cells eliciting no spikes during the 500–2000 ms period that was considered for analysis (Fig. 4*B*). Quantitatively, we computed sparsity in granule cell firing as (1 − *f*_GC_), where *f*_GC_ defined the fraction of granule cells that elicited spikes. In the default hand-tuned network, granule cells showed sparse activity, with a sparsity value of 0.79. In contrast, the interneurons showed relatively low sparsity of action potential firing in the default network (BC: 0.58; MC: 0.02; HC: 0). We asked if sparsity in GC firing was simply consequent to the divergent connectivity of 700 PP inputs to 3600 granule cells, by computing firing rates in networks where individual interneuron subtypes were removed. We deleted individual interneuron subtypes from the network by removing all synapses to and from the specific subtype, while retaining the rest of the network to match the default network. We found that deletion of either basket cells or mossy cells markedly altered firing rates of the other neuronal subtypes, with the most dramatic differences observed with the removal of basket cells from the default network (Fig. 4*B*). HIPP cells were unaffected by removal of interneurons, but removal of HIPP cells did alter firing of other cell types. This was expected from the network architecture (Fig. 1*D*) where HIPP cells did not receive inputs from other local neurons and project solely to the GCs (which is connected to the other two interneuron subtypes). Notably, consistent with previous lines of evidence on the roles of interneurons in GC sparsity (Dieni et al., 2013; Stefanelli et al., 2016; Hainmueller and Bartos, 2018; Elgueta and Bartos, 2019; Hainmueller et al., 2024), we found that removal of any of the interneuron subtypes from the default network resulted in reduction of GC sparsity (GC sparsity after BC removal: 0; GC sparsity after MC removal: 0.69; GC sparsity after HC removal: 0.71).

We used the average firing rates and spike patterns to compute the *r_β_* or the *R_β_* metric, respectively, for both the input and the output (action potentials from all GCs) patterns. We plotted output similarity *vs.* input similarity to confirm pattern separation in the default network, for the chosen set of *P*_0_ and *P*_1_input patterns and their morphs (Fig. 4*C*). To confirm that the hand-tuned DG network can perform pattern separation on different sets of randomly generated *P*_0_–*P*_1_ patterns, we repeated our analyses with 10 additional sets of patterns. These sets of morphed patterns were presented to the same default network and pattern separation performance was assessed by plotting output *vs*. input correlations (Supplementary Fig. 4A*– B*). We found that the default network manifested pattern separation with all these additional input sets as well (Supplementary Fig. 4A–B). These results confirmed that the default network was not biased to the set of patterns used to hand-tune the network but could separate additional sets of input patterns as well.

In these analyses, as GCs constitute the network output to the CA3, we computed output correlation from the action potentials of the GCs. Did the outputs of the other neuronal subtypes in the same default network manifest pattern separation with identical inputs? To answer this, we computed *r_β_* and *R_β_*for the outputs of each interneuron subtype and plotted them against the respective input correlation values. We found that the outputs of basket cells and mossy cells showed pattern separation (Supplementary Fig. S4*C–D*), but outputs of HIPP cells did not (Supplementary Fig. S4*E*). As HIPP cells receive inputs solely from PP inputs (Fig. 1*D*), the absence of pattern separation in these cell types was to be expected. In contrast, basket cells and mossy cells receive inputs from GCs (Fig. 1*D*), implying that pattern separation in these cell types could be inherited from GCs. Alternately, as they send projections back to the GCs (Fig. 1*D*), they could be playing a role in shaping pattern separation in GC outputs.

To understand how different interneurons regulated pattern separation of GC outputs in the default network, we assessed pattern separation in the models where individual interneuron subtypes were deleted. We found that the GC outputs from networks that lacked any of the three interneuron subtypes were ineffective in executing pattern separation (Fig. 4*D*), irrespective of whether pattern separation was assessed with the *r_β_* or the *R_β_* metric. The largest impact on the GC output correlations was observed when the basket cells were deleted (Fig. 4*D*). Together, these observations demonstrated that sparse firing and pattern separation in DG granule cells were strongly regulated by the different DG interneuron subtypes.

### Multiple pattern-separating networks were identified through an unbiased stochastic search algorithm

Our hand-tuned default model offers one instance of the DG network that efficaciously implemented pattern separation. The choice of one single hand-tuned model offers a single solution that is biased by the parameters that were chosen for that network. Such biases also have been shown to reflect in how the network responds when challenged with perturbations such as deleting individual components (Rathour and Narayanan, 2019; Mishra and Narayanan, 2021c; Ratliff et al., 2021; Gorur-Shandilya et al., 2022; Marder et al., 2022; Alonso et al., 2023; Marom and Marder, 2023; Schapiro and Marder, 2024; Calabrese and Marder, 2025). Therefore, we asked if the hand-tuned model was the only combination of synaptic weights in the DG network that could execute pattern separation. On the other extreme, is it possible that any random choice for these synaptic weight parameters yield efficacious pattern separation?

To address these questions, we modified the MPMOSS algorithm (Fig. 1*A*) to search for valid network models, with the parametric space spanning the different synaptic weights (Supplementary Table S8) and the objective specified to be efficacious pattern separation in the network. We generated 20,000 different random DG networks by picking 8 synaptic weight values from their respective uniform distributions, without changing any of the other network properties. To each of these 20,000 networks, we presented all 11 input patterns (*P*_0_ to *P*_1_) and obtained the outputs of the GCs for each pattern. We assessed the pattern separation capability of each network by plotting output correlation *vs.* input correlation, with correlation computed independently with the *r_β_* and *R_β_* metrics.

Given the large number of randomly generated network models, it was essential to develop quantitative metrics for evaluating the pattern separation ability of each network towards their validation (Fig. 5*A–B*). In doing this, we plotted *S_out_vs*. *S_in_*for all 11 patterns, independently with the *r_β_* or the *R_β_* correlation metric. For pattern separation to be realized, it was essential that *S_out_*be less than *S_in_*across the 9 intermediate morphed patterns (Fig. 5*A*). With reference to the *S_out_ vs*. *S_in_* plot, this translates to the need for all 9 intermediate points to fall below the 45-degrees (*S_out_* = *S_in_*) equality line of the plot. To codify this requirement into our quantitative measurements, we first rotated the *S_out_ vs*. *S_in_* plot by 45 degrees with the 45-degree line forming the transformed *x* axis (Fig. 5*B*). In this rotated plane, the region above and below the zero-line represented pattern completion (PC) and pattern separation (PS) respectively (Fig. 5*B*). We fitted a cubic polynomial to these rotated datapoints to arrive at quantifications that would maximize pattern separation and minimize pattern completion for our network outcomes.

**Figure 5:**
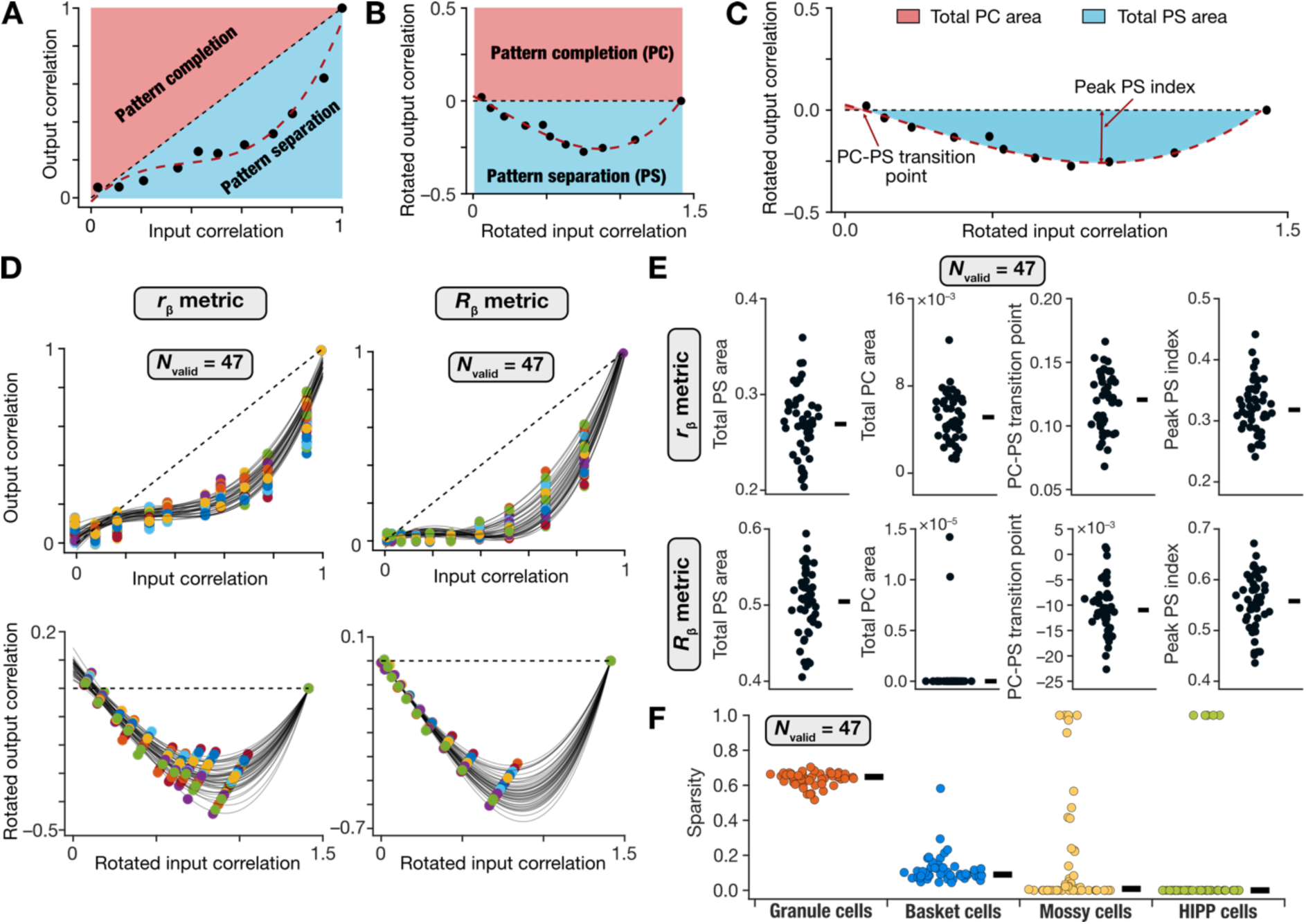
Unbiased search in the synaptic weights space yielded several DG networks that matched the performance criteria for pattern separation. (A) The *S_out_ vs*. *S_in_* plot was derived from the inputs and outputs of the network using the *r_β_* and *R_β_* metrics. The *r_β_* plot for the default network is shown for this illustration. Note that pattern separation (PS) is defined to occur if *S_out_ < S_in_*. Pattern completion (PC) occurs if *S_out_ > S_in_*. (B) The data points from the plot in panel A were rotated clockwise by 45°. A cubic polynomial was fitted to the rotated datapoints. In this rotated plane, pattern completion lies above the zero line, while pattern separation occurs below. (C) Quantitative measurements derived from the rotated plot of output *vs.* input correlation were used for validating models based on pattern separation performance. The plot and the fitted cubic polynomial shown are reproduced from panel B. Four measurements were defined using the fitted cubic polynomial: *A_PS_*, the total pattern separation area; *A_PC_*, the total pattern completion area; *T_PCPS_*, pattern completion to pattern separation transition point; and *PS_max_*, the peak pattern separation index. Effective pattern separation was deemed to be achieved if *A_PS_* > 0.15, *A_PC_* < 0.02, *A_PCPS_* < 0.25, and *PS_max_* > 0.2 (Supplementary Table S9). (D) Pattern separation plots for the 47 networks that satisfied all four measurement bounds computed with both *r_β_* and *R_β_* metrics. (E– F) Distributions of the four pattern separation measurements for *r_β_* (E; *top*) and *R_β_* (E; *bottom*) metrics as well as sparsity of different cell types (F) for all 47 valid DG networks.

We defined four quantitative measurements using the cubic polynomial fit to the *S_out_vs*. *S_in_* plot in the rotated plane (Fig. 5*C*). We defined *total pattern separation area*, *A_PS_*, as the total area under the curve of the negative-rectified cubic polynomial. *Total pattern completion area*, *A_PC_*, was computed as the total area under the curve of the positive-rectified cubic polynomial. The point at which the fitted polynomial crossed zero was defined as the *PC-PS transition point*, *T_PCPS_*. The absolute value of the peak negative deflection of the fitted cubic polynomial represented the *peak pattern separation index, PS*_max_. With these quantitative definitions, effective pattern separation was achievable by maximizing *A_PS_*, minimizing *A_PC_*, minimizing *T_PCPS_*, and maximizing *PS*_max_ (Fig. 5*C*; Supplementary Table. S9).

We imposed bounds on each of these four measurements (Supplementary Table S9) for a network to be called a pattern separation network. We computed each of these measurements from the *S_out_vs*. *S_in_*plot spanning all 11 patterns for each of the 20,000 randomly generated networks, independently with the *r_β_* or the *R_β_* correlation metric. A network was declared to be a valid pattern separation network only if all 4 measurements (*A_PS_*, *A_PC_*, *T_PCPS_*, and *PS*_max_) were within their respective bounds, with both *r_β_* and the *R_β_* correlation metrics. Thus, a total of eight measurements (four with *r_β_* and four with *R_β_*) were used for validating each network. The validation process yielded 47 valid DG networks, which was a small proportion of the generated random networks (∼0.2% of 20,000), that performed pattern separation (Fig. 5*D–E*).

### Pattern-separating networks with similar granule cell activity manifested dissimilar activity in the interneurons

How distinct were these 47 valid networks from each other in terms of the pattern separation measurements and the activity of the different neuronal subtypes? Although all valid networks satisfied all measurement bounds (for pattern separation) by requirement, we found marked network-to-network variability in their pattern separation (Fig. 5*C–E*) and sparsity (Fig. 5*F*) measurements. Importantly, although all 47 networks showed very similar sparsity values for their granule cells, the sparsity values for the different interneurons manifested pronounced network-to-network variability (Fig. 5*F*). To explore this aspect of network-to-network variability, we first computed pairwise similarity between granule cell outputs of all 47 DG network using *r_β_* and *R_β_* measures. Using these pairwise similarity measures, we identified four networks with highest similarity in terms of granule cell outputs and sparsity values (Fig. 6*A*). It may be noted that while the raster plots associated with 50 granule cells showed similar sparse firing across all four networks, action potential firing in mossy cells and basket cells were starkly different across the four networks (Fig. 6*A*). With the network architecture (Fig. 1*D*), HIPP cells did not receive any feedback from other neuronal subtypes in the network and therefore did not show marked variability across these four networks. Importantly, the pattern separation measurements of these four networks were also comparable, computed either using *r_β_* or *R_β_* measures (Fig. 6*B*). These examples illustrate our observation that networks that had very similar granule cell activity and pattern separation performance were endowed with pronounced network-to-network variability in interneuron activity.

**Figure 6:**
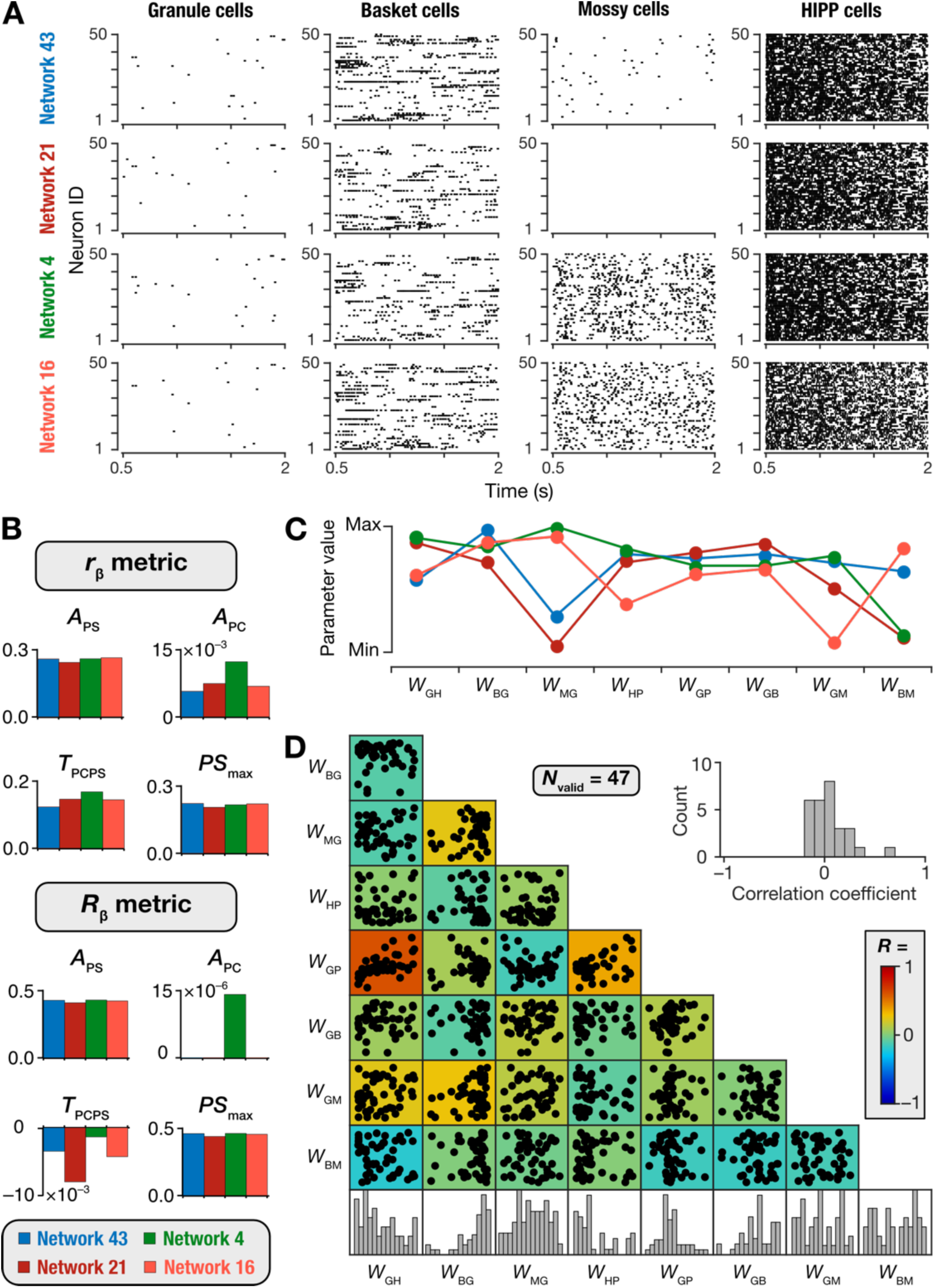
Networks with similar pattern separation performance and granule cell population activity manifested dissimilar activity across interneurons and pronounced variability in synaptic connectivity. (A) Raster plots showing firing of 50 granule (GC), basket (BC), mossy (MC), and HIPP cells (HC) of 4 selected networks where granule cell population activity was most correlated (computed through both *r_β_* and *R_β_* metrics for activity of all granule cells across networks). Although the granule cell population activity was correlated, the activity of interneurons manifested pronounced variability across the 4 networks. Sparsity values for all neurons of each subtype in the networks also followed the same trend, showing similar sparsity for GCs but widely variable sparsity for the other interneurons. Network 43: GC: 0.63, BC: 0.05, MC: 0.41, HC: 0; Network 21: GC: 0.62, BC: 0.13, MC: 1, HC: 0; Network 4: GC: 0.62, BC: 0.1, MC: 0, HC: 0; Network 16: GC: 0.6, BC: 0.06, MC: 0, HC: 0; (B) Distributions of the four pattern separation measurements for *r_β_* (*top*) and *R_β_* (*bottom*) metrics for the four selected networks. (C) Distribution of the synaptic weights for 4 selected networks. (D) Pair-wise scatter plots and distributions of the 8 synaptic weights for all 47 valid DG networks. The color-coded heat map represents Pearson’s correlation coefficient values for each pairwise relationship. The inset depicts the distribution of all the 28 unique correlation coefficients in the scatter plot matrix.

### Complexity and degeneracy in pattern-separating networks

How distinct were these 47 valid networks from each other in terms of the synaptic weights that defined them? To address this, we first considered the 4 networks that manifested similar granule cell activity (Fig. 6*A*) and pattern separation performance (Fig. 6*B*). We plotted the synaptic weights that connected the different neuronal subtypes within each of these four networks (Fig. 6*C*). We found striking variability in the synaptic connectivity across these different networks, with even extreme scenarios where some networks lacked certain connections. For instance, in Network 21, mossy cells did not elicit action potentials (Fig. 6*A*; mossy cell sparsity for Network 21 was 1) as there was weak connectivity from granule cells to mossy cells in this network (Fig. 6*C*). Thus, despite strong similarities in DG activity (Fig. 6*A*) and in pattern separation performance (Fig. 6*B*) across 4 different networks, the underlying synaptic connectivity was extremely variable (Fig. 6*C*).

To compare similarities of synaptic parameters across all 47 valid networks, we plotted the histograms of and pairwise correlations between the synaptic weight parameters governing these networks (Fig. 6*D*). We found that the synaptic weight distributions spanned a large range of the search space (histograms in Fig. 6*D*). Most synaptic weight parameters exhibited weak pairwise correlations (Fig. 6*D*), except for relatively high correlation value between the excitatory PP-to-GC and inhibitory HC-to-GC weights, which was expected due to the network architecture (Fig. 1*D*). Specifically, granule cells and HIPP cells receive common inputs from PP, and this strong correlation indicates the need to balance the excitation from the perforant pathway and the inhibition from the HIPP cells. Thus, valid pattern-separating networks did not cluster around a specific type of connectivity in achieving pattern separation through sparse firing across the granule cell population. Together, these results provide clear lines of evidence for synaptic degeneracy in heterogeneous DG network models, whereby disparate combinations of synaptic connectivity yielded similar pattern separation capabilities in DG networks.

### Network-to-network variability in the impact of deleting individual interneuron subtypes on pattern separation

Our analyses showed that deletion of individual interneuron subtypes differentially hampered firing rates, sparsity, and pattern separation performance capabilities of granule cell outputs (Fig. 4*B*, Fig. 4*D*). These observations demonstrated that sparsity and pattern separation in DG networks are not solely dependent on the projection of low-dimensional PP inputs to a high-dimensional space involving a larger number of granule cells. There are several lines of evidence for important roles for the local DG network (Fig. 1*D*) and the different interneuron subtypes in sparse firing and pattern separation (Dieni et al., 2013; Stefanelli et al., 2016; Hainmueller and Bartos, 2018; Elgueta and Bartos, 2019; Hainmueller et al., 2024). However, our observations on the differential roles of the individual neuronal subtypes were confined to the default network (Fig. 4) and are prone to biases driven by connectivity patterns in the default network. Therefore, we extended our interneuron-deletion analyses to the 47 valid pattern-separating networks, especially considering the variability in synaptic connectivity and interneuron firing properties across these networks (Fig. 5*E*, Fig. 6).

We created a total of 3 × 47 = 141 networks that were altered by 3 different perturbations (where either all BCs, all MCs, or all HCs were deleted) in each of the 47 valid networks. We presented the 11 morphed patterns to each of these networks and computed sparsity of GC firing. We quantified pattern separation performance by assessing output correlations using *r_β_* and *R_β_* across the granule cells in each of the 141 networks (Fig. 7*A–B*; Supplementary Table S10). First, we found that the impact of deletion on pattern separation was differential across different neuronal subtypes. Specifically, there was a striking loss of pattern separation performance across all networks when basket cells were deleted. However, deletion of either mossy or HIPP cells did not have as large an impact across networks (Fig. 7*A–B*). Second, and more strikingly, there was pronounced network-to-network variability in how deletion of specific interneuron subtypes affected different networks. Whereas deletion of either mossy cells or HIPP cells had a large impact on pattern separation performance in certain networks, in other networks the impact was negligible (Fig. 7*A–B*). In some of the networks, deletion of interneuron population even resulted in enhanced pattern separation capabilities, resulting in a diversity of network responses to deletions (Fig. 7*A–B*). Remarkably, however, deletion of any of the three interneuron subtypes resulted in significant loss in sparsity of granule cells, basket cells, and mossy cells (Fig. 7*C*; Supplementary Table S10).

**Figure 7:**
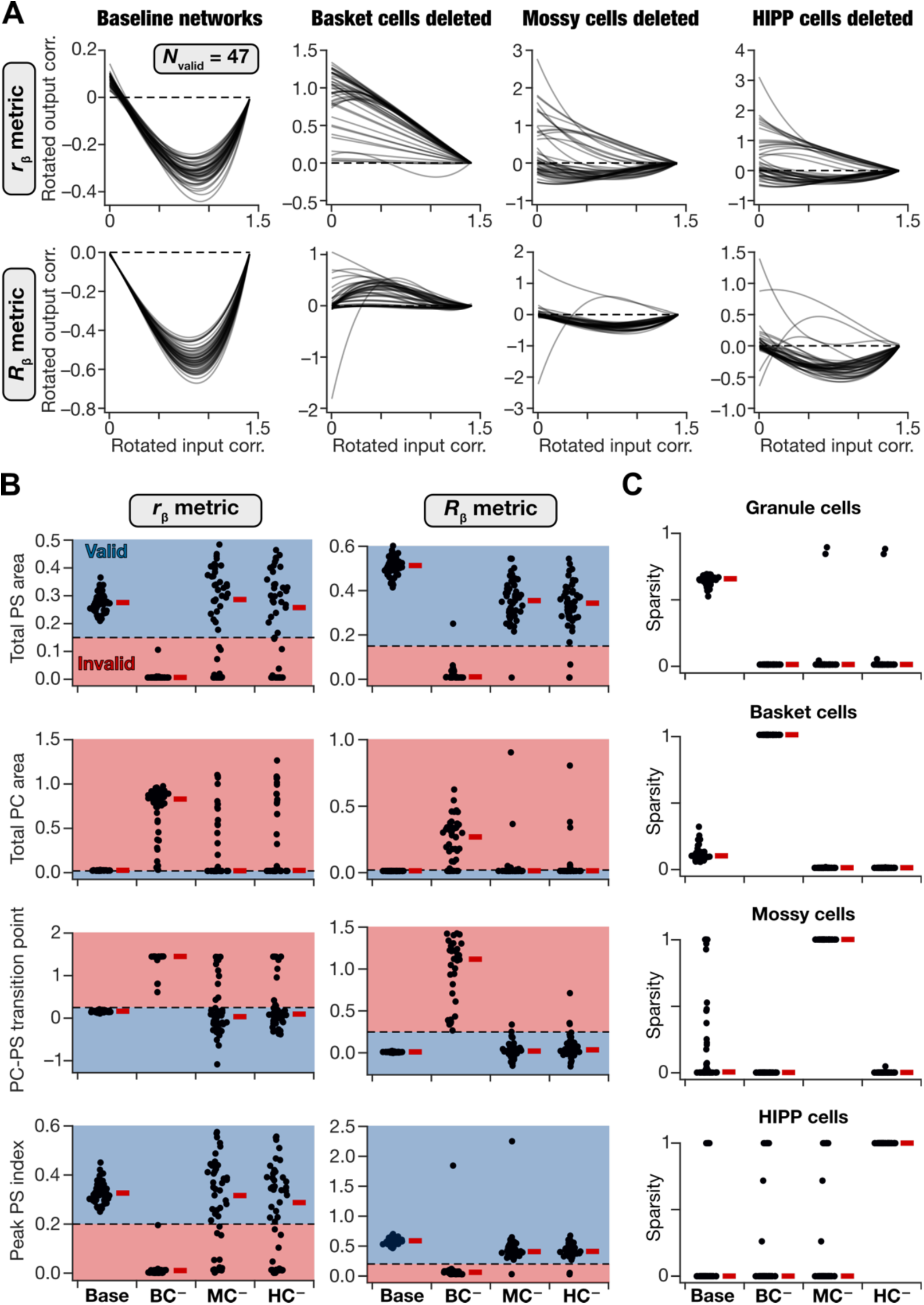
Pronounced network-to-network variability in the impact of removing individual interneuron subtypes on pattern separation performance and neural activity. (A) Visualization of pattern separation performance in all 47 valid baseline networks, and in each of the 47 networks with basket cells (BC), mossy cells (MC), or HIPP cells (HC) individually removed. Shown are fitted cubic polynomial on the rotated output correlation *vs*. rotated input correlation datapoints for all 11 morphed patterns. Correlation was computed either with the *r_β_* metric (*top*) or the *R_β_* metric (*bottom*). The pronounced variability in the impact of deletion might be observed across networks. (B) All 4 pattern separation measurements in the valid networks and in networks where individual interneuron subtypes were deleted. These measurements were calculated from the traces shown in panel A for all 47 networks, with correlation computed either with *r_β_* (*left*) and *R_β_* (*right*). Blue and red shades indicate valid and invalid ranges for each measurement, respectively. (C) Sparsity of all neuronal subtypes in all 47 valid baseline networks and after deletion of individual interneuron subtypes. In (B–C), red lines indicate the respective median values. Results of statistical analyses for these panels are presented in Supplementary Table S10.

Together, we demonstrate that the local DG circuit played a crucial role in regulating pattern separation across different networks. However, we report pronounced network-to-network variability in how interneurons affect pattern separation in the DG network, depending on the specific connectivity patterns in the network under consideration.

### Heterogeneous networks were more robust to synaptic perturbations

The high proportion of random networks that were declared invalid (∼99.8%; Fig. 6) demonstrates that pattern separation was highly sensitive to synaptic weight parameters. In biological networks, synaptic weights could undergo changes as the animals learn other tasks or undergo pathological changes. Heterogeneities in neural circuits have been proposed as a physiological mechanism to maintain resilience to perturbations in different kinds of neural circuits (Mishra and Narayanan, 2021c; Ratliff et al., 2021; Gorur-Shandilya et al., 2022; Marder et al., 2022; Rich et al., 2022; Alonso et al., 2023; Marom and Marder, 2023; Schapiro and Marder, 2024; Calabrese and Marder, 2025). Do heterogeneous DG networks maintain robust pattern separation in the face of synaptic perturbations? Would a homogeneous DG network maintain similar robust pattern separation in the presence of synaptic perturbations?

To address these questions, we compared the performance of heterogeneous and homogeneous dentate gyrus networks under different network perturbations. We introduced perturbations through jitter in synaptic weights and/or additive noise to synaptic currents. Both jitter and noise were Gaussian in nature, with increasing variances representing different graded levels. We assessed network’s pattern separation performance in all 47 homogeneous and 47 heterogeneous networks, each tested at different graded levels of jitter and/or noise (Fig. 8, Supplementary Figs. S5–S10). We computed the percentage change in all eight pattern separation measurements (*A_PS_*, *A_PC_*, *T_PCPS_*, and *PS*_max_ computed with *r_β_* and *R_β_* metrics) with reference to measurements from respective networks with no perturbations.

**Figure 8:**
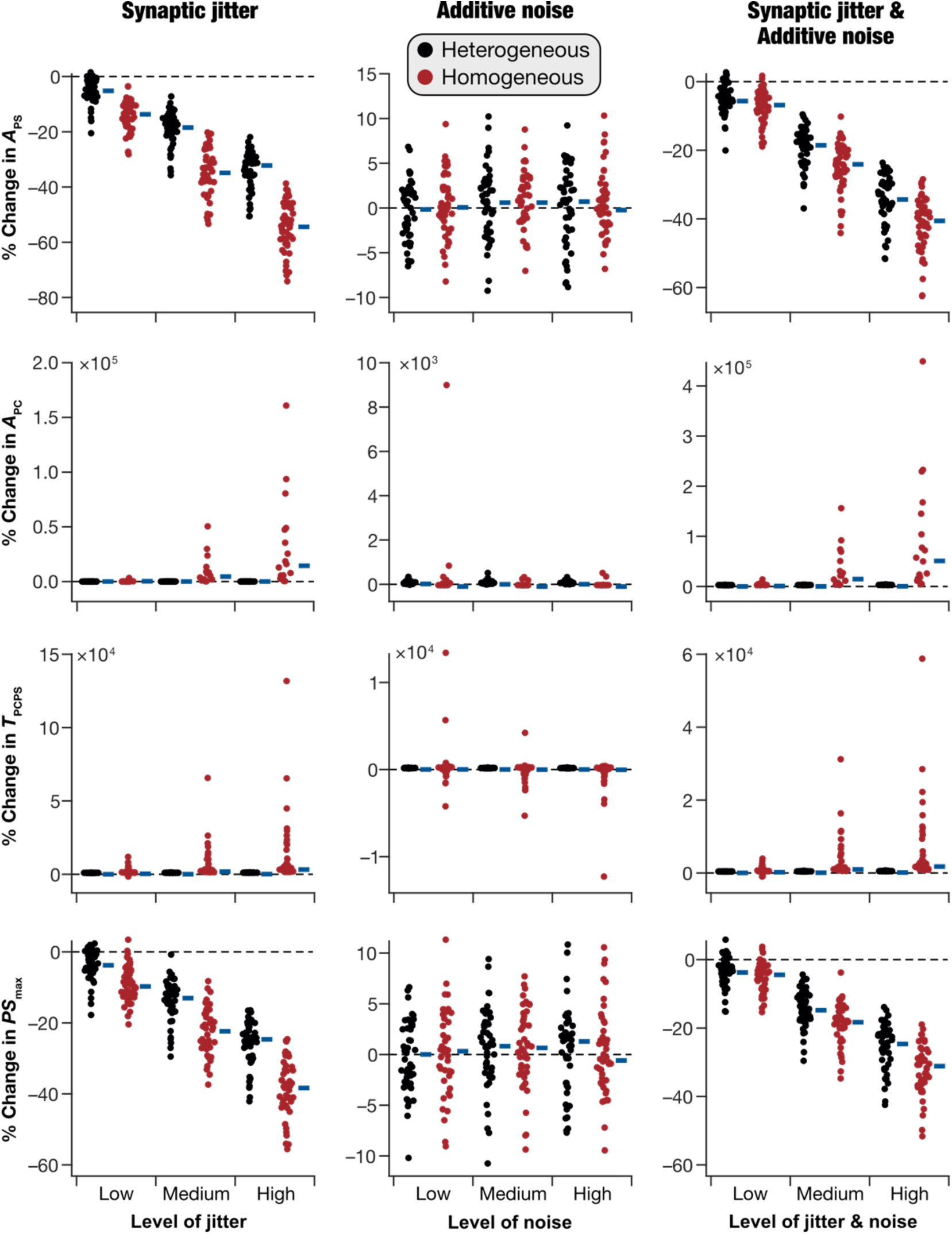
Heterogeneous dentate gyrus networks showed higher robustness in pattern separation performance when challenged with synaptic perturbations. Percentage changes in pattern separation measurements (computed using the *r_β_* metric; see Supplementary Fig. S11 for the same measurements with the *R_β_*metric instead) when perturbations of different levels were introduced. Percentage changes were computed with reference to the base networks where there were no perturbations. Perturbations were introduced in three distinct levels (low, medium, and high) as synaptic jitter (*left column*), or as additive noise to synaptic currents (*middle column*) or a combination of both jitter and noise (*right column*). The pronounced network-to-network variability, in both homogeneous and heterogeneous networks, in percentage changes of pattern separation measurements for the same levels of jitter and/or noise may be noted. In all plots, blue lines indicate the respective median values. Results of statistical analyses for these panels are presented in Supplementary Table S11.

When these metrics were computed with *r_β_* for synaptic jitter (Fig. 8; Supplementary Figs. S5–S6), we found that *A_PS_*(Fig. 8) and *PS*_max_(Fig. 8) showed significantly higher reduction (Supplementary Table S11) in homogeneous networks compared to their heterogeneous counterparts, indicating that heterogeneous networks were more robust to synaptic perturbations. Consistent with this, *A_PC_* (Fig. 8) and *T_PCPS_* (Fig. 8) showed increases in homogeneous networks with synaptic jitter but did not change in heterogeneous networks. These observations demonstrated that pattern separation performance of networks progressively degraded with graded increase in synaptic jitter, with heterogeneous networks more robust to jitter than their homogeneous counterparts. For the same set of networks, we found no significant difference in pattern separation performance between homogeneous and heterogeneous networks perturbed with synaptic jitter, when the *R_β_* metric was used for correlation computation (Supplementary Fig. S11).

We repeated these analyses with additive noise (Fig. 8; Supplementary Figs. S7–S8) to synaptic currents as well as with the combination of jitter and noise (Fig. 8; Supplementary Figs. S9–S10). We did not observe significant changes in pattern separation performance (computed either with *r_β_*or *R_β_*metrics) of homogeneous *vs*. heterogeneous networks when additive noise was introduced into synaptic currents (Fig. 8; Supplementary Figs. S9–S10; Supplementary Table S11). We observed significant differences in robustness between heterogeneous and homogeneous networks when perturbations in both synaptic jitter and additive noise were introduced together, with the *r_β_* metric (Fig. 8; Supplementary Figs. S9– S10; Supplementary Table S11) but not with the *R_β_* metric (Supplementary Figs. S9–S11).

## DISCUSSION

How do we explain the several conflicting lines of evidence on the roles of different local-circuit interneurons of the dentate gyrus to pattern separation and sparsity? In addressing this question, we built model populations that matched biological heterogeneities in the functional characteristics of the four major DG neuronal subtypes using unbiased stochastic search algorithms. We constructed heterogeneous DG networks with non-repeating units of each neuronal subtype, with neuronal proportions, connectivity, and synaptic receptor properties adopted from their biological counterparts. We presented these networks with progressively morphed patterns as inputs and plotted network output similarity *vs.* input similarity. We chose two different correlation-based measures, one on average firing rates and another on spike trains, to compute similarity among input or output patterns. We generated a large population of random DG networks through unbiased sampling of the synaptic weights space and connected the heterogeneous populations of different neuronal subtypes with these synaptic weights. For each of these random networks, we computed network responses to morphed inputs and calculated output similarity values for each of the several morphed input pairs. We found a small subset of randomly generated DG networks where output similarity was less than the respective input similarity values across all morphed patterns, which we declared as pattern-separating networks.

We found that these valid networks showed effective pattern separation performance and similar sparsity in granule cell firing. Strikingly, however, we found pronounced network-to-network variability in firing rates and sparsity of interneuron populations, in synaptic weight values that connected different neuronal subtypes, and in how deletion of interneuron populations affected pattern separation and sparsity. Thus, disparate combinations of non-random and non-unique synaptic connectivity yielded similar pattern separation capabilities in heterogeneous DG networks, offering clear evidence for synaptic degeneracy and complexity in the manifestation of pattern separation in DG networks. Finally, synaptic perturbation analyses demonstrated that pattern separation performance (measured with correlation of average firing rates) showed progressive degradation with increasing level of jitter in synaptic weights, but not with additive noise. Remarkably, we found heterogeneous DG networks to be more resilient to synaptic jitter compared to their homogeneous counterparts. The pronounced nature of network-to-network variability and the beneficial roles of intrinsic heterogeneities in imparting resilience present strong arguments for systematic characterization and analyses of different forms of heterogeneities in studying pattern separation in the DG. These analyses also demonstrate that pattern separation in heterogeneous DG networks emerge through intricately woven, non-random, and non-unique synaptic interactions among its several neuronal components.

### Complexity and multi-scale degeneracy in heterogeneous pattern-separating DG networks

Degeneracy, the ability of disparate structural components to yield similar functional outcomes, is a ubiquitous feature of biological systems across all scales of analysis (Edelman and Gally, 2001; Price and Friston, 2002; Mason, 2010; Cropper et al., 2016; Rathour and Narayanan, 2019; Goaillard and Marder, 2021; Mishra and Narayanan, 2021a; Seenivasan and Narayanan, 2022; Albantakis et al., 2024; Mittal and Narayanan, 2024; Calabrese and Marder, 2025). Our analyses demonstrate the manifestation of multi-scale degeneracy in pattern separating DG networks. At the cellular scale, consistent with prior studies from different DG neuronal subtypes (Mishra and Narayanan, 2019; Mishra and Narayanan, 2021c, b; Shridhar et al., 2022; Schneider et al., 2023; Kumari and Narayanan, 2024), we found that disparate parametric combinations yielded signature functional characteristics of individual neurons belonging to four different subtypes (Fig. 1; Supplementary Fig. S1). At the network scale, several non-unique combinations of synaptic weight combinations yielded pattern separating DG networks with sparse firing (Fig. 5–6). The concurrent expression of cellular and network-scale degeneracy emphasizes the inherent flexibility and adaptability of the DG network (Mishra and Narayanan, 2016; Mishra, 2019; Mishra and Narayanan, 2019; Mishra and Narayanan, 2021c), with several degrees of freedom available towards achieving pattern separation.

The manifestation of heterogeneities and degeneracy rules out the presence of a unique, completely determined solution that yields functionally precise neuronal models (at the cellular scale) or a pattern-separating DG network (at the network scale). In addition, we found large proportions of randomly generated neuronal models (Fig. 1*B*) or network models (∼99.8%; 19,953 of 20,000) to be invalid. Therefore, in both cellular and network scales, arbitrarily random combinations couldn’t yield collective functional outcomes. These observations provide direct evidence that pattern-separating DG networks are neither fully determined nor are completely random. They require specific non-random interactions among components to yield collective neuronal or circuit function. The manifestation of such intermediate levels of randomness along with the expression of degeneracy indicate that pattern-separating networks are complex systems (Watts and Strogatz, 1998; Barabasi and Albert, 1999; Edelman and Gally, 2001; Kim and Wilhelm, 2008; Alon, 2019; Adler and Medzhitov, 2022; Alon, 2023).

Complex systems execute their collective function through non-unique and non-random interactions among their functionally specialized subsystems. The ability of disparate functional subsystems to yield similar collective function through non-random, motif-based interactions is a hallmark of complex systems. Thus, our analyses argue for approaching pattern-separating DG networks from the complex systems framework. This implies that the relationship between structural components and collective functional outcomes is not one-to-one, but many-to-many.

### Multiple subsystems differentially contribute to pattern separation in heterogeneous DG networks

Our observations with the population-of-networks approach are in striking contrast with the classical Marr-Albus theory, where pattern separation is postulated to be realized by divergent feedforward excitation (Marr, 1969; Albus, 1971; Marr, 1971). First, despite being endowed with the same divergent afferent connectivity and despite being built of the same set of neurons, we found that pattern separation couldn’t be achieved in a vast majority of networks where local connectivity was randomized (Fig. 5–6). These observations demonstrate that divergent activity and low excitability of granule cells are not sufficient to guarantee pattern separation in DG networks. Instead, consistent with other observations from the DG network (Dieni et al., 2013; Stefanelli et al., 2016; Hainmueller and Bartos, 2018; Elgueta and Bartos, 2019; Hainmueller et al., 2024), our results suggest a critical role for local interneurons in regulating pattern separation and sparse firing. These conclusions are further emphasized by the strong dependency of pattern separation and sparse firing on local synaptic weights. Strong dependencies are inferred from observations that a majority of randomly connected networks were invalid (Fig. 5–6) and that deletion of interneurons (Fig. 7) or synaptic jitter (Fig. 8) degraded pattern separation performance of the networks.

Second, within the valid pattern-separating networks, we found pronounced network-to-network variability in firing rates and sparsity of interneuron populations, in local synaptic weights, and in how different interneuron populations contributed to pattern separation and sparsity (Figs. 5–6). This is consistent with the manifestation of degeneracy, where different neural circuits performing the same function (through different combinations of subsystems) show differential dependencies on individual subsystems (Mishra and Narayanan, 2021c; Ratliff et al., 2021; Gorur-Shandilya et al., 2022; Marder et al., 2022; Alonso et al., 2023; Marom and Marder, 2023; Schapiro and Marder, 2024; Calabrese and Marder, 2025). More importantly, the complex system framework and network-to-network variability provide perfect substrates for explaining discrepancies in the literature on interneuron contributions. Specifically, the several recurrent connections between excitatory and inhibitory interneurons in the DG network has yielded puzzling conclusions about the role of each neuronal subtype in regulating sparsity and pattern separation (Ratzliff et al., 2004; Jinde et al., 2012; Dieni et al., 2013; Stefanelli et al., 2016; Chavlis et al., 2017; Danielson et al., 2017; GoodSmith et al., 2017; Bui et al., 2018; Hainmueller and Bartos, 2018; Cayco-Gajic and Silver, 2019; Elgueta and Bartos, 2019; GoodSmith et al., 2019; GoodSmith et al., 2022; Hainmueller et al., 2024; Scharfman, 2025). Our observations show that there are several non-random synaptic connectivity patterns that can yield effective pattern separation, with pronounced network-to-network variability in the roles of interneuron subtypes.

Future studies should approach pattern separation in the DG network not exclusively from the perspective of one-to-one relationship between individual components and function. Instead, the focus must be on the global structure associated with how different components interact with each other in several non-unique, non-random ways to yield pattern separation. Such analyses should account for the manifestation of network-to-network variability in dependencies on specific components, ensuring that the heterogeneities across networks are respected and all networks are not lumped into one homogeneous population. It is also important to account for heterogeneities in the proportions of neuronal subtypes in different parts of the dentate gyrus (Gaarskjaer, 1978; Seress and Pokorny, 1981; Ribak and Seress, 1983; Amaral et al., 2007).

Our conclusions on degeneracy, complexity, and network-to-network variability were possible only because we used a population-of-networks approach to study the DG network. The choice of one hand-tuned model offers a single solution that is biased by the parameters that were chosen for that network. Such biases have been demonstrated to reflect in how the network responds when challenged with perturbations such as deleting a component (Rathour and Narayanan, 2014; Mukunda and Narayanan, 2017; Basak and Narayanan, 2018; Haddad and Marder, 2018; Basak and Narayanan, 2020; Gorur-Shandilya et al., 2020; Jain and Narayanan, 2020; Mishra and Narayanan, 2021c; Ratliff et al., 2021; Gorur-Shandilya et al., 2022; Mittal and Narayanan, 2022; Roy and Narayanan, 2023; Srikanth and Narayanan, 2023; Schapiro and Marder, 2024). Together, our analyses strongly advocate the use of a population-of-networks approach, involving networks with different heterogeneities, in assessing pattern separation in DG networks. The complex systems framework and the population-of-networks approach offer an ideal route to account for the several DG circuit components and heterogeneities therein, towards implementing robust pattern separation through flexible mechanistic routes.

### Heterogeneities, metrics for circuit performance, and robustness to perturbations

Biological systems are characterized by their robust execution of function despite the expression of perturbations. There are lines of evidence that the manifestation of heterogeneities could offer a biological mechanism to yield robustness in certain types of networks (Mishra and Narayanan, 2021c; Ratliff et al., 2021; Gorur-Shandilya et al., 2022; Marder et al., 2022; Rich et al., 2022; Alonso et al., 2023; Marom and Marder, 2023; Schapiro and Marder, 2024; Calabrese and Marder, 2025). Our analyses also demonstrate that DG networks endowed with intra-subtype heterogeneities were more robust to the synaptic weight perturbations compared to their homogeneous counterparts (Fig. 8). However, our analyses also highlight the need for the use of different metrics in assessing circuit function. Specifically, our analyses demonstrated that pattern separation performance computed from correlations of average firing rates (*r_β_*) showed progressive degradation with increasing level of jitter in synaptic weights, but not with additive noise. These analyses also indicated that heterogeneous networks were more resilient to synaptic jitter compared to their homogeneous counterparts (Fig. 8). In striking contrast, when we computed pattern separation performance through correlations across spike trains (*R_β_*) instead of average firing rates, pattern separation performance did not manifest large differences and there were no differences between heterogeneous and homogeneous networks as well (Supplementary Fig. S11). These analyses provide lines of evidence that the additional information carried by the temporal pattern of spikes might offer efficacious pattern separation despite perturbations (Madar et al., 2019; Bird et al., 2024). These observations also emphasize the need to use multiple metrics to assess network performance, to avoid biases associated with the use of a single metric in evaluating circuit function.

### Future directions within the complex adaptive systems framework

The complex adaptive systems perspective for the emergence of circuit function provides a unified framework to understand heterogeneities and how they contribute to collective function of the network and robustness therein. As our analyses showed, the focus here must be on how different forms of heterogeneities (*e.g.*, in neuronal intrinsic properties, local synaptic connectivity, sparsity of afferent connectivity, and neuronal morphology) interact with each other towards achieving collective function and resilience therein. Context-dependent multi-component plasticity during behavioral learning provides the means to align different forms of heterogeneities towards yielding robust function (Bliss and Gardner-Medwin, 1973; Bliss and Lomo, 1973; Greenstein et al., 1988; Pavlides et al., 1988; Shors and Dryver, 1994; Wang et al., 1997; Davis et al., 2004; McHugh et al., 2007; Lopez-Rojas et al., 2016; Luna et al., 2019; Mishra and Narayanan, 2021a, 2022). Such multi-component plasticity and its specificity to behavioral contexts could be mediated by temporally aligned adult neurogenesis, which offers a hyperplastic substrate to regulate sparsity of representation by implementing plasticity heterogeneity (Schmidt-Hieber et al., 2004; Ge et al., 2007; Dieni et al., 2013; Aimone et al., 2014; Abrous and Wojtowicz, 2015; Li et al., 2017; Lodge and Bischofberger, 2019; Mishra and Narayanan, 2019; Denoth-Lippuner and Jessberger, 2021; Huckleberry and Shansky, 2021; Mishra and Narayanan, 2021c; Shridhar et al., 2022). Structured multi-component plasticity driven by behavioral context, with adult neurogenesis as the substrate, offers a flexible set of adaptive mechanisms that recruit different circuit components to yield one of several robust solutions for sparse firing and pattern separation in DG circuits (Mishra and Narayanan, 2016; Mishra, 2019; Mishra and Narayanan, 2019; Mishra and Narayanan, 2021c, a).

Thus, the complex adaptive systems framework unifies disparate lines of research on the dentate gyrus on different forms of plasticity, adult neurogenesis, engram formation, and resource allocation towards selection of specific combinations of heterogeneities that yield robust circuit function (Schmidt-Hieber et al., 2004; Ge et al., 2007; Dieni et al., 2013; Aimone et al., 2014; Yiu et al., 2014; Park et al., 2016; Josselyn and Frankland, 2018; Lodge and Bischofberger, 2019; Mishra, 2019; Mishra and Narayanan, 2019; Pignatelli et al., 2019; Josselyn and Tonegawa, 2020; Lau et al., 2020; Huckleberry and Shansky, 2021; Mishra and Narayanan, 2021a, 2022; Shridhar et al., 2022). Importantly, findings on representational drift in network representations for the same behavioral context (Ziv et al., 2013; Driscoll et al., 2022; Keinath et al., 2022; Snoo et al., 2023) could be effectively fit within the complex adaptive systems framework, as representational drift simply implies that the complex network is traversing across different degenerate solutions that retain functional precision.

Future computational models could focus on *adaptive emergence of* effective pattern separating networks through multi-component plasticity in random networks that are endowed with different cell types, plasticity in neuronal properties and connectivity, and hyperplasticity in a subset of adult-born neurons. In pattern separating networks arrived through such an *adaptive learning* process, the manifestation of specific network motifs (Luo, 2021) that resemble connectivity in the DG microcircuit could be assessed. The specific roles of intrinsic and synaptic plasticity (across different cell types) in the emergence of pattern separating networks and underlying motifs could then be assessed within the complex adaptive systems framework. Heterogeneities across different neuronal populations and their connectivity, the contributions of temporally-aligned adult neurogenesis in encoding specific contexts, representational drift, and network-to-network variability in the role of different interneurons could be easily assessed within these emergent networks for comparison with their biological counterparts. More elaborate models could also incorporate non-repeating morphologically realistic neuronal models of each cell type, receiving specifically patterned afferent and local inputs to different parts of their dendritic arbor, other DG cell types including neuron-glia interactions, and feedback connectivity from CA3 towards exploring the contributions of these additional details to complexity and degeneracy.

Experimentally, it is important that studies that assess the roles of the DG in pattern separation systematically account for the different forms of heterogeneities that the DG is endowed with as well as animal-to-animal variability in different components that contribute to pattern separation. Importantly, analyses of neurological disorders affecting hippocampal function should consider different routes to dysfunction, including possibilities where unstructured changes to heterogeneities could result in loss of collective functions under pathological conditions (Mishra and Narayanan, 2021a; Rich et al., 2022; Stober et al., 2023). Together, the complex adaptive systems framework offers a unified framework for tying together the disparate lines of research on the heterogeneous dentate gyrus network and could be effectively harnessed for studying its multifarious functions under both health and disease conditions.

## METHODS

Neural circuits are heterogeneous. Heterogeneities span structural characteristics, intrinsic properties, local synaptic connectivity, and afferent connectivity onto individual neurons in the network. A fundamental question in neuroscience is to understand the origins, the prevalence, and the implications of these heterogeneities to neural circuit physiology. Although the pronounced manifestation of heterogeneities in different cellular-scale properties is well-characterized in the dentate gyrus (DG), the implications for these heterogeneities to its circuit-scale function have not been thoroughly assessed. Our goal here was to study the implications of different neural heterogeneities on pattern separation in the hippocampal dentate gyrus network. In addressing this, we took a population-of-models approach that involved a multi-scale cascade of unbiased stochastic searches. Specifically, four independent stochastic searches were first used to find physiologically validated heterogeneous cellular-scale models of four different neuronal subtypes. A second level of unbiased stochastic search was performed to identify networks composed of these validated neuronal models that were structurally constrained by the DG architecture and functionally validated for their ability to perform pattern separation. We used the population of networks that emerged from the second search to assess heterogeneities and degeneracy in these networks towards the emergence of pattern separation capabilities.

### Subtype-specific single-neuron models

We first ensured that the constituent single-neuron model populations functionally matched the DG neuronal subtypes and their specific heterogeneities. The DG network was constructed with four neuronal subtypes: the principal granule cells and three different interneuron subtypes, namely basket cells, mossy cells, and hilar perforant path-associated (HIPP) cells. Irrespective of neuronal subtype, we modeled individual cells using the adaptive exponential integrate-and-fire (aEIF) spiking neuronal model (Brette and Gerstner, 2005), given their ability to match different physiological characteristics and the low computational complexity. The choice of the aEIF model for implementing all subtypes allowed for exploration of intra- and inter-subtypes heterogeneities within the same parametric and measurement spaces.

The membrane potential dynamics of aEIF neurons was defined by the following system of ordinary differential equations (Brette and Gerstner, 2005):

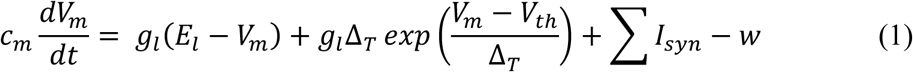

where *V_m_* represented the membrane potential (in mV), *c_m_* and *g_l_* defined the membrane capacitance (in µF) and the leak conductance (in mS), respectively. *E_l_* was the reversal potential for the leak conductance (in mV), Δ_*T*_ was the slope factor (in mV), *V_th_* was the threshold (in mV), *I_syn_* was the synaptic current (in nA). The dynamics of the adaptation parameter *w* in Eq. (1) evolved as:

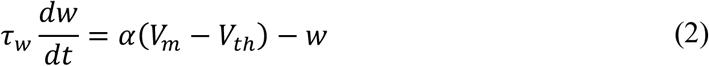

where *τ_w_* defined the time constant association with *w* and *α* was the adaptive coupling parameter. We used surface area (*A*) as representation for the structural properties of the neurons, which was used to compute *c_m_* and *g_l_* in Eq. 1 as follows:

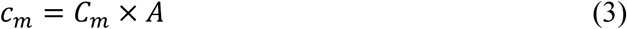

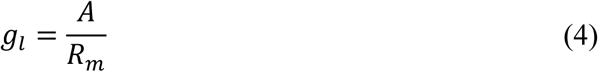

where *C_m_* and *R_m_* represented the specific membrane capacitance (in µF/cm^2^) and specific membrane resistance (in kΩ · cm^2^), respectively.

As the dynamics evolved for different synaptic inputs, the neuron was considered to have elicited an action potential every time the value of *V_m_* (computed from Eq. 1) crossed the threshold voltage *V_th_*. The occurrence of a spike also triggered updates in the values of *V_m_* and *w* as follows:

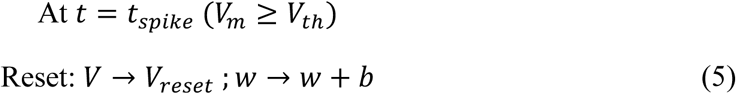

where *b* defined the spike-triggered adaptation parameter and *V_restet_* denoted the voltage to which *V_m_* was reset after a spike. A refractory period of 5 ms was imposed following every spike, during which no further spiking was allowed.

The default values of the 10 parameters that defined models for each of the four cell types are listed in Supplementary Tables S1–S4.

### Intrinsic electrophysiological properties of the four neuronal subtypes

We validated models for each neuronal subtype against six electrophysiological measurements: membrane time constant (*τ_m_*), sag ratio (Sag), input resistance (*R_in_*), spike frequency adaptation (*SFA*), and action potential firing frequencies for 50 pA and 150 pA pulse current injections (*f*_50_ and *f*_150_, respectively). All measurements were obtained after an initial delay period of 500 ms, during which the transient membrane dynamics of the model stabilized to its resting potential. These measurements were computed using established protocols for computing each of them (Lübke et al., 1998; Bartos et al., 2001; Ratzliff et al., 2002; Krueppel et al., 2011; Mishra and Narayanan, 2019) and are elaborated below.

The membrane time constant (*τ_m_*) was estimated by fitting a single exponential to the membrane potential response to a 20-pA pulse current injected into the neuron for 1500 ms:

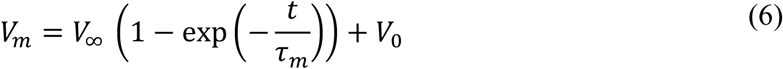

where *V*_∞_ denoted the steady-state voltage response and *V*_0_ represented the resting potential at which the pulse current was injected.

Sag ratio was measured as the ratio of the steady-state voltage deflection (*V_SS_*) to the peak voltage deflection (*V_peak_*) obtained in response to –50 pA current pulse for one second. Input resistance (*R_in_*) was calculated by injecting a family of nine 1-second-long pulse currents with amplitudes ranging from –40 pA to 40 pA in steps of 10 pA. Steady-state voltage deflections were calculated for each pulse current injection. *R_in_* was then computed as the slope of the steady-state voltage deflection *vs.* current injected plot for all 9 pulse current amplitudes. As HIPP cells elicit action potentials for current amplitudes above 20 pA, we used step currents ranging from –20 pA to 20 pA in steps of 5 pA for *R_in_*computation in HIPP cells. Firing frequencies (*f*_50_ and *f*_150_) were computed as the number of spikes elicited by the neuron upon injection of 50 pA and 150 pA pulse currents, respectively, for 1 second duration each. Spike frequency adaptation (SFA) was measured as the ratio of the first inter-spike interval to the last inter-spike interval for a pulse current of 150 pA amplitude injected for 1 second. The electrophysiological ranges for all six measurements derived from previous electrophysiological studies (Lübke et al., 1998; Bartos et al., 2001; Ratzliff et al., 2002; Krueppel et al., 2011; Mishra and Narayanan, 2019) are listed in Supplementary Table S5.

### Generation of heterogeneous populations of functionally validated models for the four neuronal subtypes

We needed physiologically validated heterogeneous neurons of all four neuronal subtypes to construct the DG network. We used four independent implementations of multi-parametric multi-objective stochastic search (MPMOSS) algorithm (Fig. 1*A*) to generate heterogeneous model populations of all four neuronal subtypes. The MPMOSS algorithm has been widely used to study the emergence of signature physiological characteristics in neurons and their networks through disparate combinations of different parameters (Foster et al., 1993; Prinz et al., 2003; Taylor et al., 2009; Marder and Taylor, 2011; Rathour and Narayanan, 2012; Rathour and Narayanan, 2014; Sinha and Narayanan, 2022), including in DG neurons with different model complexities (Mishra and Narayanan, 2019; Mishra and Narayanan, 2021c; Shridhar et al., 2022; Schneider et al., 2023; Kumari and Narayanan, 2024). We chose an unbiased random search approach covering a large swath of the parametric space, rather than confining search spaces to constrictive subspaces, to allow for model heterogeneities to potentially span the entire parametric space.

In implementing MPMOSS, we first generated one valid model for each neuronal subtype by hand-tuning the parameters such that all 6 measurements were in their respective valid range (Supplementary Table S5). These models then served as the substrate for performing the stochastic search where we specified upper and lower bounds for each parameter around the parameters of the default network (Supplementary Table S1–S4). To ensure that the search remained unbiased to any specific distribution, these bounds were used to define uniform distributions for each parameter for each neuronal subtype. Each parameter was randomly sampled several times (*N* = 100,000 for granule cells, 60,000 for basket cells, 10,000 for mossy cells, and 20,000 for HIPP cells) from its respective distribution, together creating several multi-parametric random models of each neuronal subtype (Fig. 1*A*).

All six electrophysiological measurements were computed for each of these random models and were independently validated against the electrophysiological measurement ranges of the respective neuronal subtype (Supplementary Table S5). A model was declared valid only if all measurements were within their respective physiological ranges; otherwise, the model was considered invalid. This validation process, applied to every generated model across the four neuronal subtypes, yielded a subset of models that were valid (*N_valid_*) for each neuronal subtype. The heterogeneities within and across valid models of different neuronal subtypes, in the 9-dimensional parametric space and the 6-dimensional measurement space, were visualized using principal component analysis. In addition, statistical analyses involving the histogram of individual parameters and pairwise correlations between valid model parameters/measurements were performed for each of the four neuronal subtypes to assess heterogeneities.

### Dentate gyrus network model endowed with within-subtype heterogeneities

Valid models of the four neuronal subtypes were then used to construct the heterogeneous DG network with non-repeating units. We built a network made of 3600 granule cells, 500 basket cells, 180 mossy cells, and 50 HIPP cells (Fig. 1*D*). The relative proportions of each cell type were informed by previous studies (Andersen et al., 2006; Myers and Scharfman, 2009; Chavlis et al., 2017; Mishra and Narayanan, 2019). The primary afferent input to the DG is the perforant pathway (PP), which we modeled using 700 independent spike trains, with specific firing rates and spike timing distributions. Granule cells function as the primary output neurons of the DG, projecting excitatory synapses onto CA3 pyramidal cells.

The connectivity within the DG circuitry (Fig. 1*D*) was adopted from previous work (Chavlis et al., 2017). Specifically, HIPP cells, as GABAergic interneurons, established inhibitory synapses onto granule cells. Granule cells made excitatory synaptic connections with mossy cells and basket cells. Mossy cells, which are glutamatergic interneurons, connected to both granule and basket cells through excitatory synapses. Basket cells, which are inhibitory interneurons, established inhibitory connections with granule cells. The afferent inputs (PP) and all the neuronal subtypes together formed 8 types of synapses in the model. The specific connectivity between individual neurons or with afferent inputs was established randomly, by specifically accounting for the different connection probabilities (Supplementary Table S6) derived from an earlier study (Chavlis et al., 2017). For the default network, the weights of all connections were hand-tuned towards achieving effective pattern separation by the DG network (see below).

### Modeling synapses

Given the nature of the individual neuronal subtypes that were used to construct the network, there were excitatory (glutamatergic) and inhibitory (GABAergic) synapses that connected different neurons in the network. The excitatory synapses were modelled as synaptic current mediated through colocalized AMPA and NMDA receptors, which were activated upon spiking of the respective presynaptic neuron. The inhibitory synapses were modelled as current mediated through GABA_A_ receptors. The currents through AMPA and GABA_A_ receptors were modelled using the following formulation:

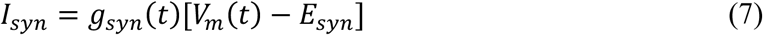

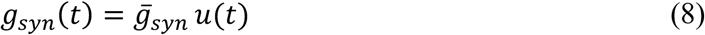

where, *I_syn_* represented the synaptic current, *E_syn_* denoted the reversal potential of the receptor (0 mV for AMPA and NMDA receptors and −90 mV for GABA_A_ receptors), *V_m_* defined the membrane potential and *g_syn_*(*t*) tracked the evolution of receptor conductance. *g_syn_*(*t*) was computed as a scaler multiple of *u*(*t*), a double exponential waveform to represent the rise and decay of synaptic currents.

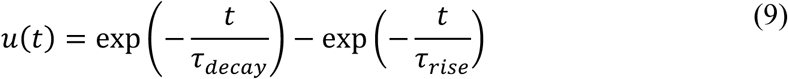

where *τ_rise_* and *τ_decay_* defined the time constants associated with the rise and decay of synaptic conductance, respectively. The values for *τ_rise_* and *τ_decay_* for the receptors associated with the different synapses in the network (Supplementary Table S7) were taken from previous studies (Geiger et al., 1997; Bartos et al., 2001; Larimer and Strowbridge, 2008; Myers and Scharfman, 2009; Chavlis et al., 2017). *g̅_syn_* defined the peak conductance of the AMPA and GABA_A_ receptors and will be referred to as the weight of the connection.

The dependence of the current through NMDA receptor on postsynaptic membrane voltage was modelled as:

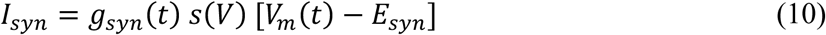

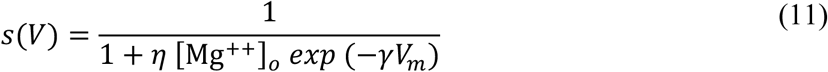

where *s*(*V*) defined the additional magnesium-dependent mechanism of the NMDA receptors. The parameters that defined the sigmoid, *η* was set at 0.28 mM^−1^, [*Mg*^++^]_*o*_ was 1 mM, and *γ* was 0.04 mV^−1^. Synaptic conductance *g_syn_*(*t*) for NMDARs followed a similar formulation as in Eq. (8–9). *g̅_syn_* of NMDA receptors was defined as a scaled version of the *g̅_syn_* of AMPA receptors (Supplementary Table S7), with a fixed NMDAR:AMPAR ratio used for scaling (Chavlis et al., 2017).

We introduced delays between presynaptic spike and the onset of the corresponding postsynaptic responses. Different values for delays were used for different connections (Supplementary Table S7), which were taken from previous studies (Geiger et al., 1997; Bartos et al., 2001; Larimer and Strowbridge, 2008; Myers and Scharfman, 2009).

### Modeling perforant pathway inputs

Our model contained 700 PP inputs impinging on the granule and HIPP cells. We modelled 700 PP inputs as 700 independent spike trains generated from a Poisson distribution with *λ* = 8 Hz and inter-spike interval generated from an exponential distribution with *λ* = 125 ms 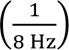. We generated two distinct input patterns *P*_0_ and *P*_1_ by randomly generating 700 spike trains for each pattern. These two patterns formed the extreme inputs that were fed as inputs to the DG network. We confirmed the distinctiveness of these two patterns by ensuring that the input correlation measures computed between the two patterns approached zero. To assess pattern separation capabilities of the network, we used the morphed input approach where *P*_0_ was progressively morphed into *P*_1_ through several intermediate input patterns (Leutgeb et al., 2007; Renno-Costa et al., 2010; Mishra and Narayanan, 2016; Mishra, 2019). Specifically, we generated 9 more intermediate input patterns by progressively morphing *P*_0_ to *P*_1_, yielding a total of 11 input patterns *P*_0_, *P*_0.1_, *P*_0.2_, …, *P*_0.9_ and *P*_1_, referred to as *P_β_*with *β* representing the morphing parameter. With increasing *β*, patterns progressively became more dissimilar to *P*_0_and more alike *P*_1_. For generating pattern *P_β_*, *β* fraction of the 700 PP inputs were randomly picked from the *P*_1_ pattern and remaining (1 − *β*) fraction of PP inputs were picked from pattern *P*_0_. We presented each of these 11 input patterns to the network and obtained the outputs of the granule cells in the network for each input pattern. We quantified pattern separation by plotting the similarity/dissimilarity among the granule cell outputs *vs*. the similarity/dissimilarity among the input patterns that were presented to elicit those outputs. This was repeated for each of the 11 patterns to cover a range of input similarities. Pattern separation was deemed to be implemented by the network if the similarity among the network outputs was lower than the similarity among the network inputs (Leutgeb et al., 2007; Renno-Costa et al., 2010). All patterns were 2 seconds in duration. We allowed the network to stabilize for the first 500 ms and considered the remaining 1500 ms for computing similarity metrics.

### Similarity metrics

We rigorously tested six different metrics, used in previous studies (Mishra, 2019; Mishra and Narayanan, 2019; Bird et al., 2024; Wang et al., 2024), to identify those that satisfied our three criteria: invariance to population size, average firing rate across the population, and the specific form of the firing rate distribution of input patterns. For all the metrics defined below, *N* denoted the number of neurons in the population, *F_β,i_* represented the normalized firing rate of *i*^th^ neuron of the *P_β_* pattern. Hamming distance (*HD_β_*) between *P*_0_ and *P_β_* was computed as:

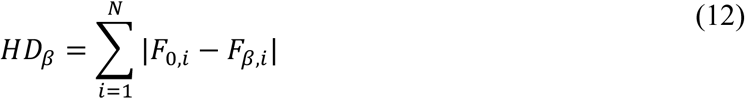

Normalized Euclidean Distance (*nED_β_*) between *P*_0_ and *P_β_* was computed as:

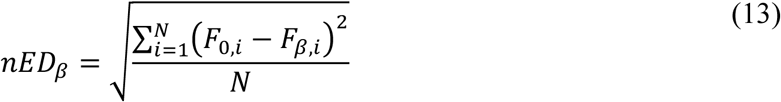

Cosine similarity (*θ_β_*) between *P*_0_ and *P_β_* was computed as:

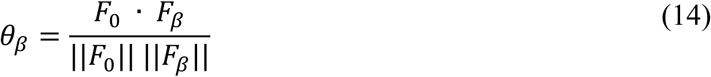

where *F_β_* represented the vector of average firing rate of *P_β_*, *F*_0_ · *F_β_* denoted the dot product between *F*_0_ and *F_β_*, and ||*F_β_*|| defined the *L*_2_ norm of vector *F_β_*.

Mutual information (*MI_β_*) between *P*_0_ and *P_β_* was computed as:

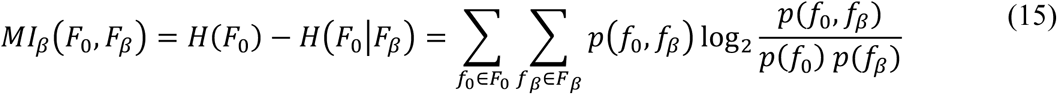

where *F_β_* defined the vector of average firing rate of *P_β_*, *p*(*f_β_*) represented the marginal probability of firing rates in pattern *P_β_*, and *p*(*f*_0_, *f_β_*) denoted the joint probability of firing rates in patterns *P*_0_ and *P_β_*.

The correlation metric between average firing rates (*r_β_*) between *P*_0_ and *P_β_* was defined as:

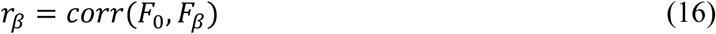

where *corr*() represented the Pearson correlation, and *F_β_* defined the vector built of the average firing rate of each neuron in pattern *P_β_*.

To compute the correlation metric between instantaneous firing rates (*R_β_*) between *P*_0_ and *P_β_*, we first computed the instantaneous firing rate of all the neurons when these two patterns were presented as network inputs. This was accomplished by convolving the spike train with a Gaussian function with *μ* = 0 and 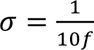, where *f* was the average firing rate of the spike train. This resulted in *N* rows corresponding to the instantaneous firing rates of *N* neurons in each pattern (Supplementary Fig. S3). We then computed pairwise Pearson’s correlations between the instantaneous firing rates of all *N* neurons for each of the two patterns under considerations, *P*_0_ and *P_β_*. We computed *R_β_* as the Pearson’s correlation between the resulting *N* × *N* similarity matrices *C*_0_ and *C_β_*, associated respectively with *P*_0_and *P_β_*.

To compare the dependence of these 6 similarity/dissimilarity metrics on the size of the population, we generated three independent sets of *P*_0_and *P*_1_. The population size of the three sets of generated inputs were 1000, 2000, and 3000. We morphed each set of *P*_0_–*P*_1_ pairs to generate 11 patterns for each set (3×11 patterns) and calculated all 6 metrics for 3 set of morphed patterns. To assess the invariance of these metrics to average firing rates, we repeated a similar process involving 3 sets of *P*_0_–*P*_1_ pairs with average firing rates of 5 Hz, 15 Hz, and 25 Hz. We considered Poisson *vs*. exponentially distributed firing frequencies to address the question of whether these metrics were invariant to the specific form of firing rate distribution. For each of the three factors (population size, average firing rate, and distribution type), we generated 10 independent sets of patterns and analyzed whether the mean similarity/dissimilarity varied across different conditions.

### Unbiased stochastic search in synaptic weights space to identify heterogeneous networks performing effective pattern separation

We generated one DG network capable of executing pattern separation by hand-tuning the eight synaptic weights. However, as hand tuning yields a single biased network that does not account for network-to-network variability in synaptic connectivity, we resorted to searching for DG networks that were effective in performing pattern separation across the synaptic weight space. We modified the unbiased stochastic search (MPMOSS) strategy that we had used to identify valid single-neuron models to now search for valid network models that were effective in performing pattern separation. The parametric search space was defined by the 8 synaptic weights between the different cell types, which were picked randomly from respective uniform distributions for the underlying conductances (Supplementary Table S8). We generated 20,000 random networks by sampling the synaptic weights space and validated them against bounds on measurements that defined effective pattern separation (Fig. 5*B*; Supplementary Table S9). The population-of-networks approach involving unbiased random search spanning multiple parameters and multiple validation measurements allowed us to explore variability across different valid networks that performed pattern separation.

### Assessing the impact of individual interneuron subtypes on pattern separation in the population of DG networks

To assess the role of the specific interneuron subtypes (*i.e.*, basket cells, mossy cells, and HIPP cells) in pattern separation performance by the heterogeneous DG networks, we systematically deleted each interneuron population from the network and computed pattern separation metrics. For deletion of individual neuronal subtypes, we set the weights of all the synapses formed onto or by that interneuron subtype to zero. We repeated this procedure for each of the 3 interneuron subtypes for all *N_valid_* number of valid DG networks (total 3 × *N_valid_* networks).

We plotted *S_out_ vs*. *S_in_* spanning all 11 patterns for each of the 3 × *N_valid_* DG networks, using both the *r_β_* and *R_β_* metrics. We computed the different pattern separation measurements for all these networks and compared these measurements with their base counterparts where all interneurons were intact.

### Assessing robustness of the pattern separation performance in the heterogeneous and homogeneous DG network population to synaptic jitter and noise

We compared pattern separation performance of heterogeneous and homogeneous DG networks when perturbations are introduced into the synaptic weights and/or synaptic currents. We used the *N_valid_* number of heterogeneous networks generated from the stochastic search to generate a population of homogeneous networks. We implemented this by replacing the non-repeating population of neurons in the heterogeneous networks by a homogeneous set of base neuron models. In other words, in the homogeneous network all GCs were identical to each other and so were each of the 3 different interneuron subtypes (one per subtype). This gave us one homogeneous network corresponding to each of the *N_valid_* heterogeneous networks, with conserved synaptic connectivity between the homogeneous and heterogeneous counterparts.

We introduced perturbations to the network either as jitter to the synaptic weights of the network or as additive noise to the synaptic current or both. For synaptic weight perturbation, we introduced random jitter to each synaptic weight, picked from a Gaussian distribution with mean zero and standard deviation *σ*. We introduced three levels of jitter: low (*σ* = 1), medium (*σ* = 3), and high (*σ* = 5). We introduced additive zero-mean Gaussian noise to the synaptic current with three levels: low (*σ* = 1), medium (*σ* = 3), and high low (*σ* = 5). When both jitter and noise were present together, we perturbed both the synaptic weight and added additive Gaussian noise, with the same three levels of perturbations. We computed the different pattern separation metrics using both the *r_β_* and *R_β_* metrics for all *N_valid_*heterogeneous networks and their homogeneous counterparts to compare pattern separation performance between homogeneous *vs*. heterogeneous networks.

### Computational details

All simulations and analyses were performed within the MATLAB programming environment (Mathworks Inc.) with custom written codes. The integration time step (Δ*t*) was fixed at 0.1 ms for all the simulations. All statistical analyses were performed using R (R Core Team, 2013).

## ACKNOWLEDGMENTS

The authors thank Dr. Poonam Mishra and members of the cellular neurophysiology laboratory for helpful discussions and for comments on a draft of this manuscript. This work was supported by the Wellcome Trust-DBT India Alliance (Senior fellowship to R. N.; IA/S/16/2/502727) and the Ministry of education (R. N. & S. S.).

