## Supplementary Figures S1-S11, Supplementary Tables S1-S11 for "Degeneracy explains diversity in interneuronal regulation of pattern separation in heterogeneous dentate gyrus networks"

### Supplementary material

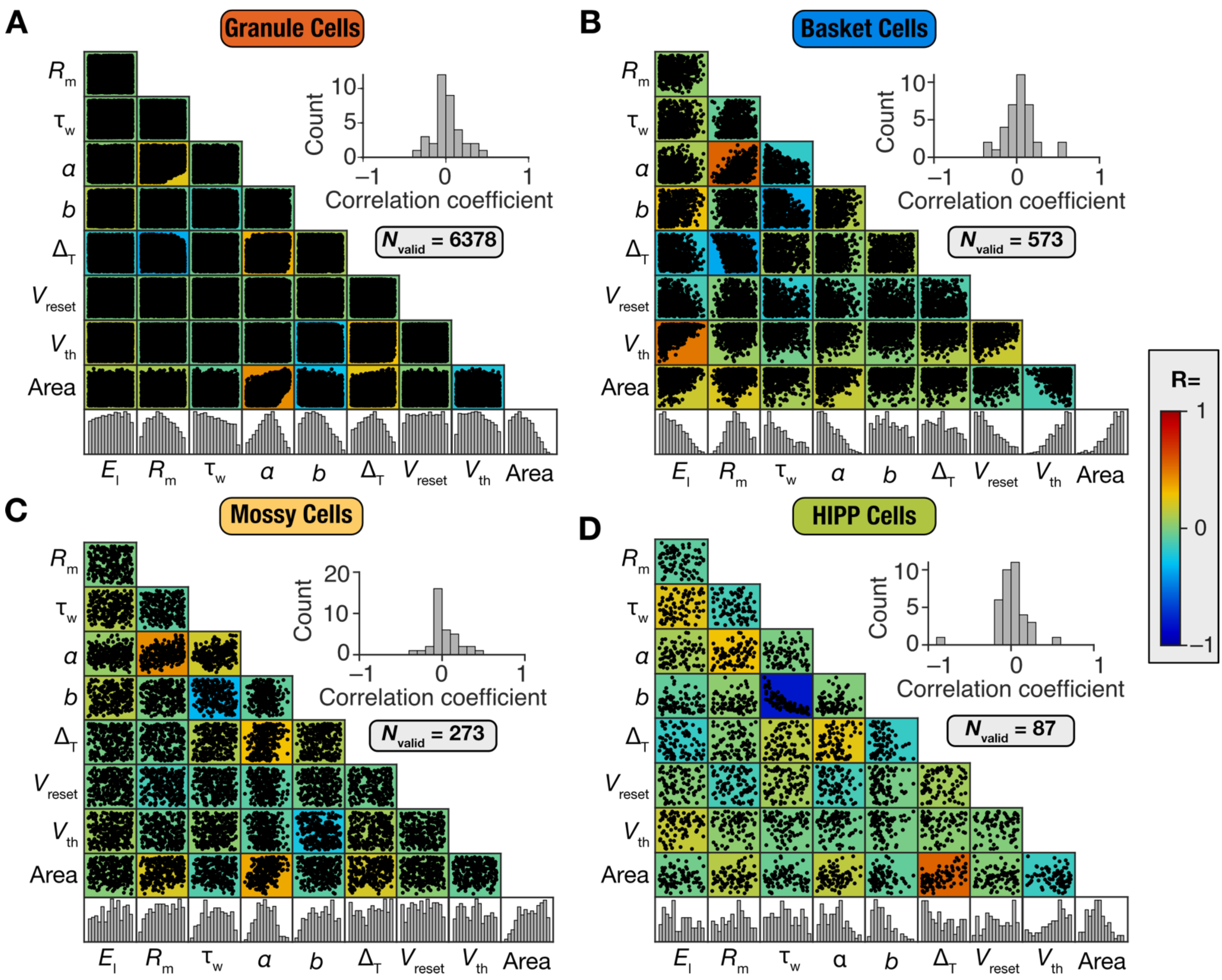

**Supplementary Figure S1: Weak pairwise correlations among heterogeneously distributed parameters of model populations of four neuronal subtypes.** (A–D) Scatter plot matrix of all parameters from valid populations of granule (A), basket (B), mossy (C), and HIPP (D) cells obtained using MPMOSS. The bottom most rows in each panel indicates the histogram of the specific parameter, showing widespread heterogeneity across models. The insets within each panel depict the histogram of the Pearson's correlation coefficient values, which are overlaid on the respective scatter plot matrices, for the unique correlation coefficient values from the pairwise comparisons shown in the scatter plot matrix. The capacitance value was uniformly set at 1 $\mu F/cm^2$ across all models and therefore is not included in these plots.

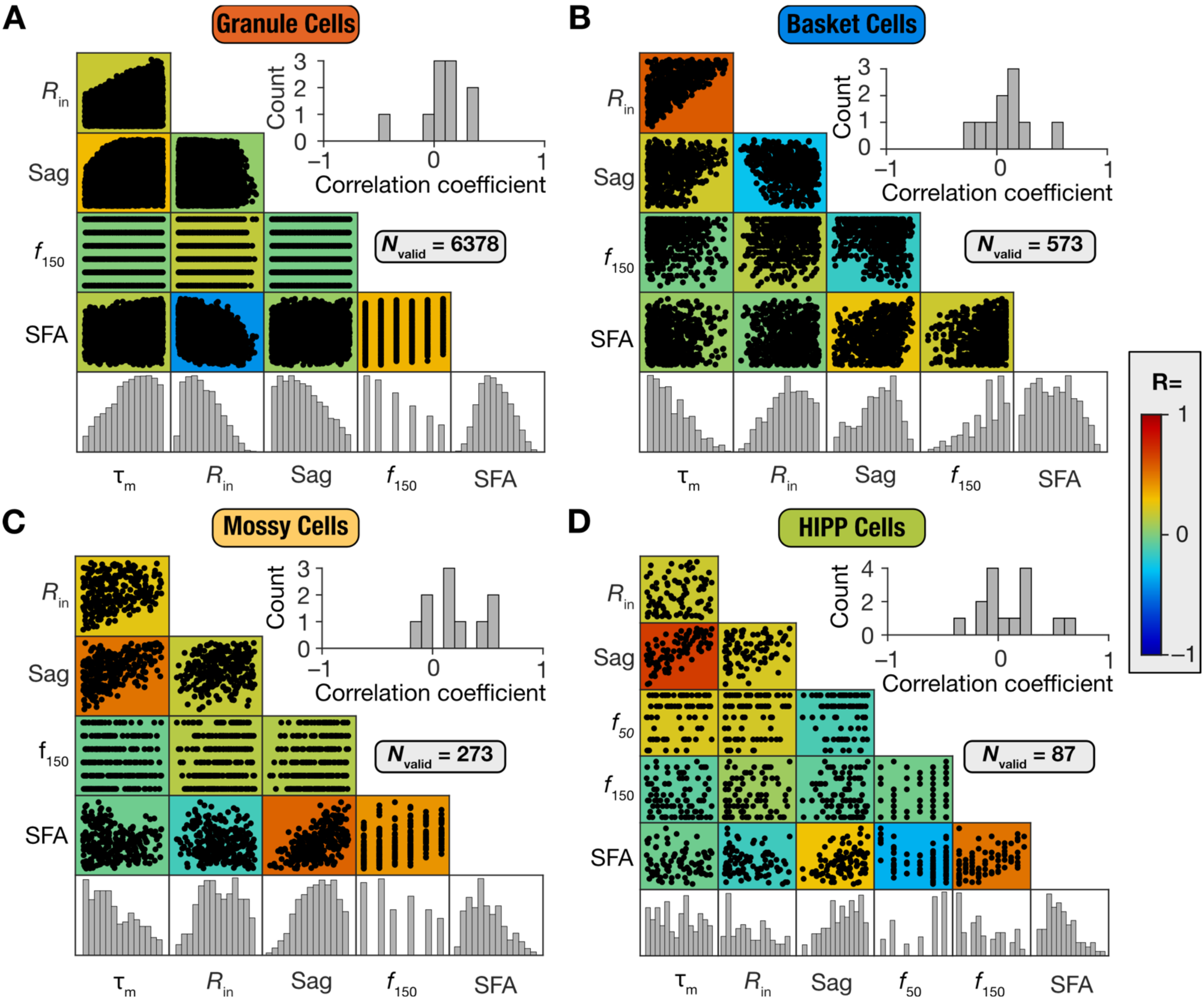

**Supplementary Figure S2: Weak pairwise correlations among heterogeneously distributed measurements of model populations of four neuronal subtypes.** (A–D) Scatter plot matrix of measurements from valid populations of granule (A), basket (B), mossy (C), and HIPP (D) cells obtained using MPMOSS. The bottom most rows in each panel indicates the histogram of the specific measurement, showing widespread heterogeneity across models. The insets within each panel depict the histogram of the Pearson's correlation coefficient values, which are overlaid on the respective scatter plot matrices, for the unique correlation coefficient values from the pairwise comparisons shown in the scatter plot matrix. The firing rate for 50 pA current injection, $f_{50}$, was required to be identically zero across all valid GC, BC, and MC models and therefore are not part of these scatter plots.

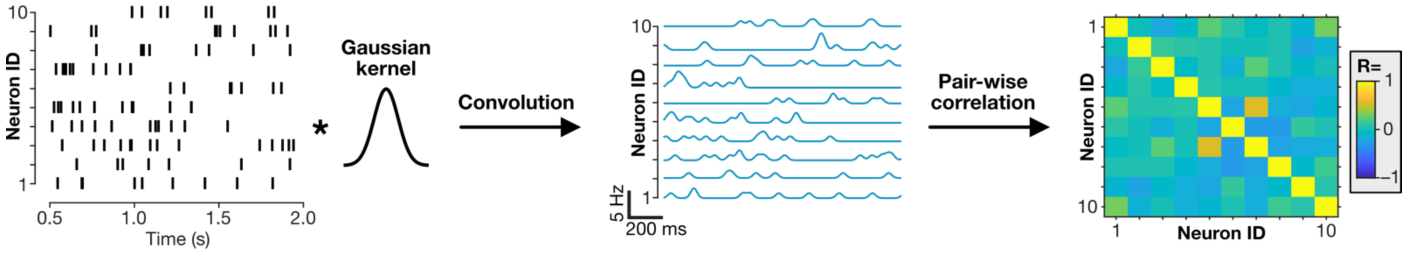

**Supplementary Figure S3: Quantification of pattern separation using correlation in instantaneous firing rates ($R_\beta$).** For calculation of $R_\beta$ between two patterns, spike times of both the patterns were first convolved with a Gaussian kernel of $\sigma = \frac{1}{10f}$, where $f$ represented the mean firing rate of the population. Then pairwise correlations were computed between each neuron pair, which gave us an $N \times N$ matrix for each pattern. $R_\beta$ between any two patterns was computed as the correlation coefficient between the matrices associated with the two patterns.

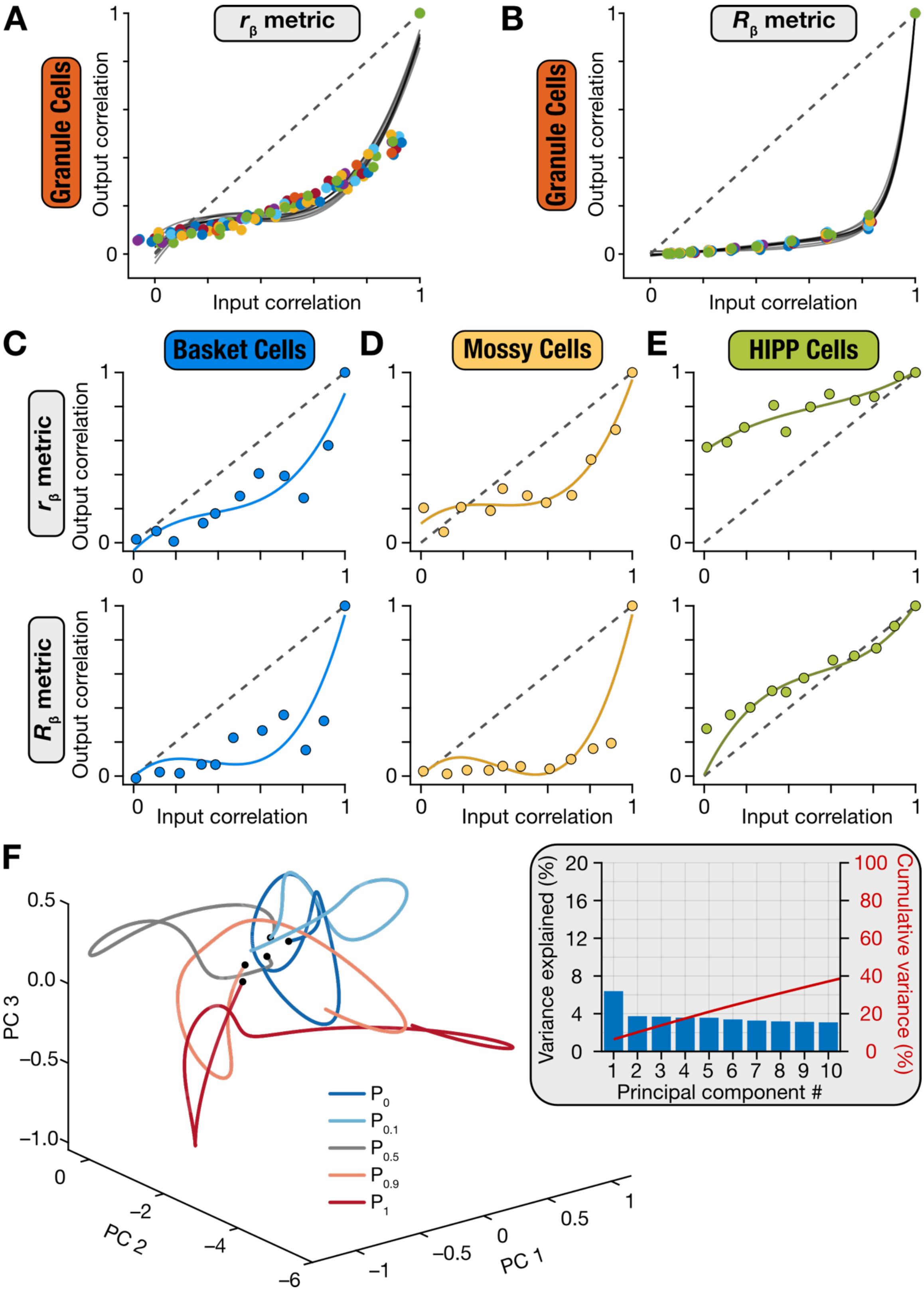

**Supplementary Figure S4: Pattern separation performance and dynamics of the default hand-tuned DG network.** (A–B) The hand-tuned DG network was tested with 10 distinct randomly generated input pattern sets. Pattern separation performance was comparable across all patterns, irrespective of whether correlation was computed used quantified using $r_\beta$ (A) or $R_\beta$ (B). These analyses confirmed that the network's pattern separation ability was not restricted to the default input set (shown in Fig. 4). (C–E) Pattern decorrelation plots for the basket cell (C), mossy cell (D), and HIPP cell (E) populations quantified using $r_\beta$ (*top*) or $R_\beta$ (*bottom*) metrics for the default input set. (F) The temporal evolution of instantaneous firing rates of the 3600 granule cell population for 5 different input patterns plotted in a lower-dimensional space using PCA. The black dot marks the start of the trajectory for each case. *Inset*, The variance explained by the first 10 principal components plotted individually and as a cumulative sum. Note the low variance explained by the first 10 dimensions for the sparse and decorrelating DG network.

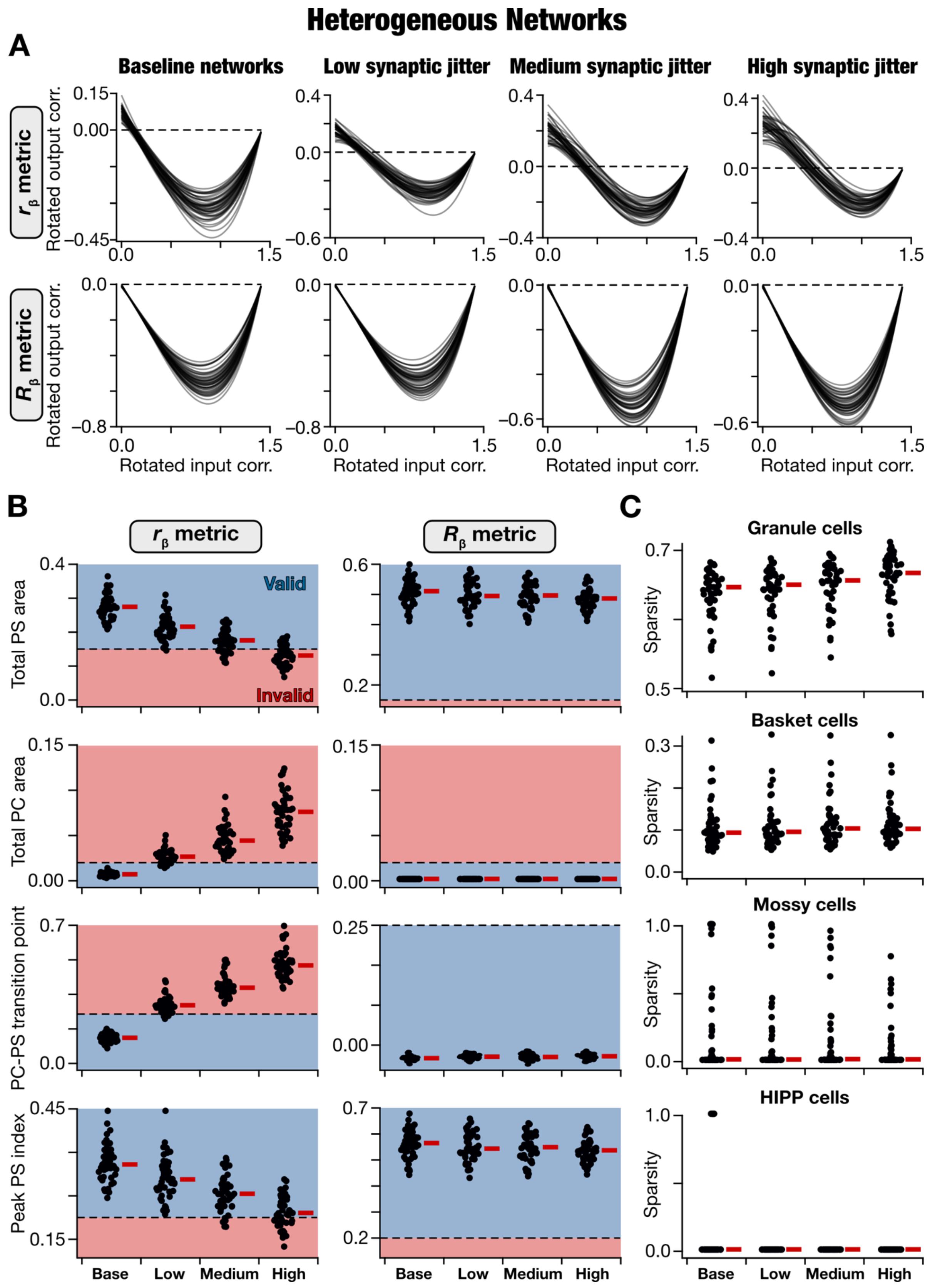

**Supplementary Figure S5: Network-to-network variability in the impact of introducing synaptic jitter on pattern separation performance of the 47 valid heterogeneous DG networks.** (A) Visualization of pattern separation performance in all 47 valid baseline heterogeneous networks, and in each of the 47 networks with low, medium, and high levels of synaptic jitter. Shown are fitted cubic polynomial on the rotated output correlation *vs*. rotated input correlation datapoints for all 11 morphed patterns. Correlation was computed either with the $r_\beta$ metric (*top*) or the $R_\beta$ metric (*bottom*). The pronounced variability in the impact of jitter might be observed across networks. (B) All 4 pattern separation measurements in the valid networks and in networks where different levels of synaptic jitter was introduced. These measurements were calculated from the traces shown in panel A for all 47 networks, with correlation computed either with $r_\beta$ (*left*) and $R_\beta$ (*right*). (C) Sparsity of all neuronal subtypes in all 47 valid baseline networks and after introduction of synaptic jitter of different levels. In (B–C), red lines indicate the respective median values.

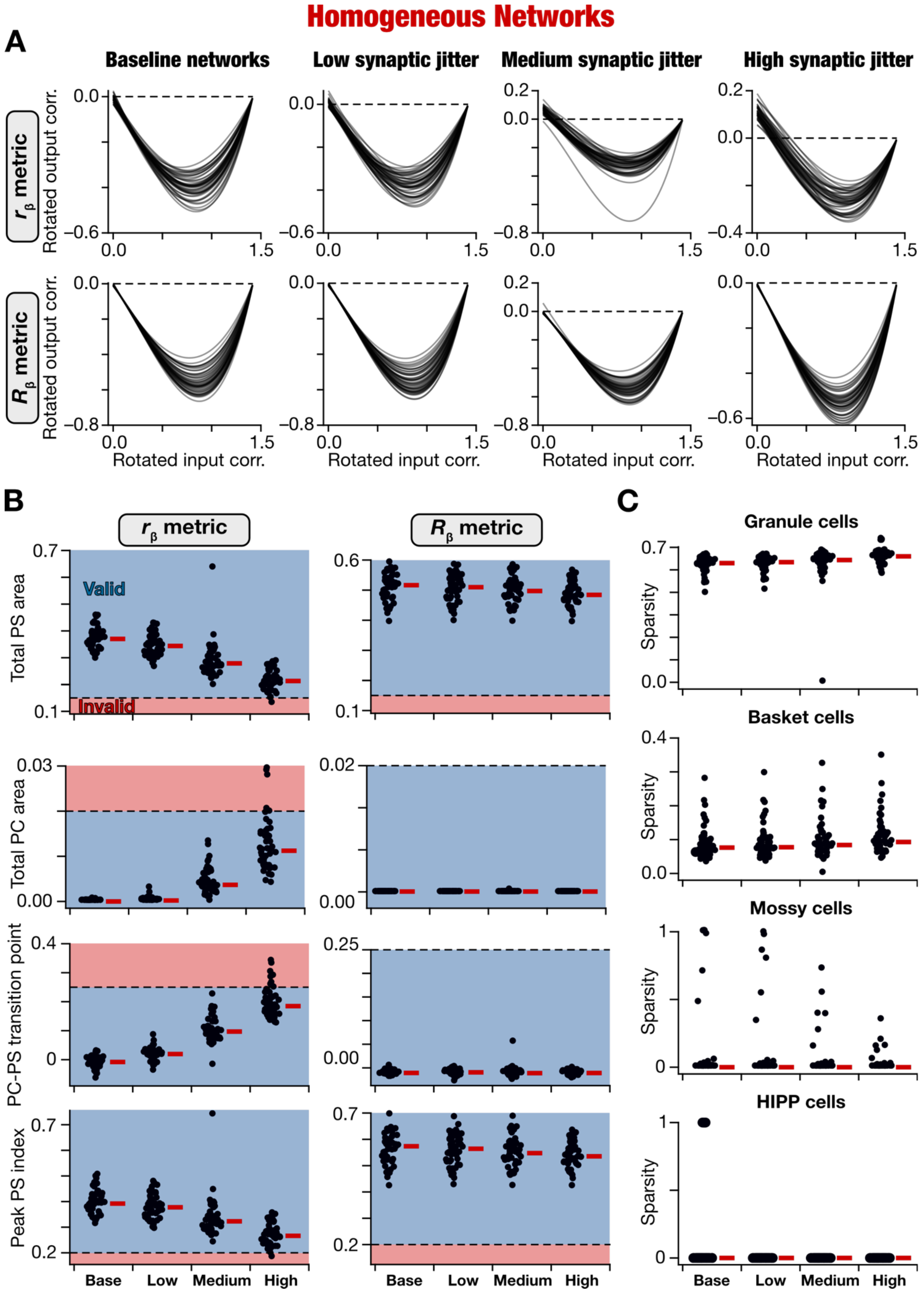

**Supplementary Figure S6: Network-to-network variability in the impact of introducing synaptic jitter on pattern separation performance of the 47 valid homogeneous DG networks.** (A) Visualization of pattern separation performance in all 47 valid baseline homogeneous networks, and in each of the 47 networks with low, medium, and high levels of synaptic jitter. Shown are fitted cubic polynomial on the rotated output correlation *vs*. rotated input correlation datapoints for all 11 morphed patterns. Correlation was computed either with the $r_\beta$ metric (*top*) or the $R_\beta$ metric (*bottom*). The pronounced variability in the impact of jitter might be observed across networks. (B) All 4 pattern separation measurements in the valid networks and in networks where different levels of synaptic jitter was introduced. These measurements were calculated from the traces shown in panel A for all 47 networks, with correlation computed either with $r_\beta$ (*left*) and $R_\beta$ (*right*). (C) Sparsity of all neuronal subtypes in all 47 valid baseline networks and after introduction of synaptic jitter of different levels. In (B–C), red lines indicate the respective median values.

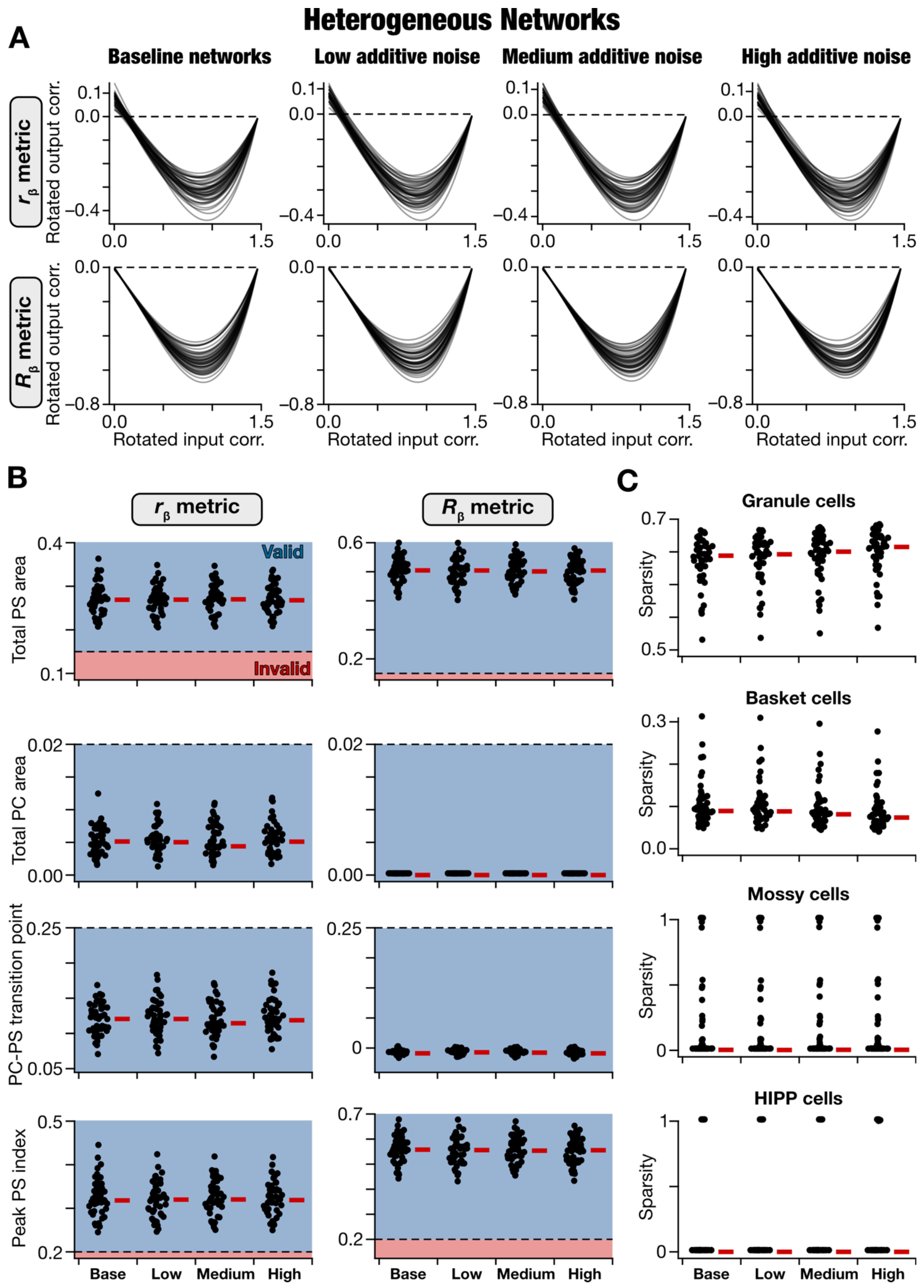

**Supplementary Figure S7: Network-to-network variability in the impact of introducing additive noise on pattern separation performance of the 47 valid heterogeneous DG networks.** (A) Visualization of pattern separation performance in all 47 valid baseline heterogeneous networks, and in each of the 47 networks with low, medium, and high levels of additive noise. Shown are fitted cubic polynomial on the rotated output correlation *vs*. rotated input correlation datapoints for all 11 morphed patterns. Correlation was computed either with the $r_\beta$ metric (*top*) or the $R_\beta$ metric (*bottom*). (B) All 4 pattern separation measurements in the valid networks and in networks where different levels of additive noise was introduced. These measurements were calculated from the traces shown in panel A for all 47 networks, with correlation computed either with $r_\beta$ (*left*) and $R_\beta$ (*right*). (C) Sparsity of all neuronal subtypes in all 47 valid baseline networks and after introduction of additive noise of different levels. In (B–C), red lines indicate the respective median values.

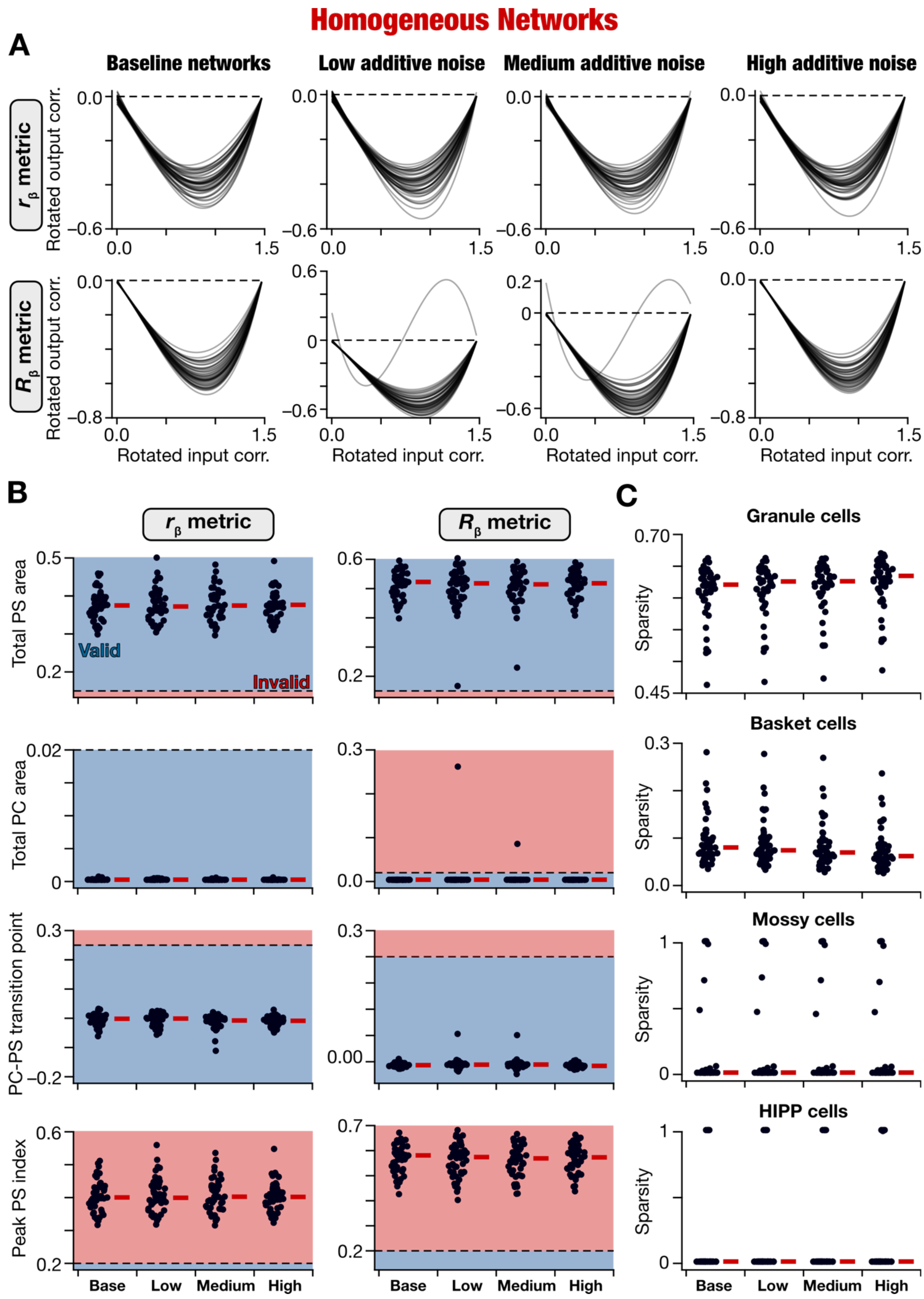

**Supplementary Figure S8: Network-to-network variability in the impact of introducing additive noise on pattern separation performance of the 47 valid homogeneous DG networks.** (A) Visualization of pattern separation performance in all 47 valid baseline homogeneous networks, and in each of the 47 networks with low, medium, and high levels of additive noise. Shown are fitted cubic polynomial on the rotated output correlation *vs*. rotated input correlation datapoints for all 11 morphed patterns. Correlation was computed either with the $r_\beta$ metric (*top*) or the $R_\beta$ metric (*bottom*). The pronounced variability in the impact of noise might be observed across networks. (B) All 4 pattern separation measurements in the valid networks and in networks where different levels of additive noise was introduced. These measurements were calculated from the traces shown in panel A for all 47 networks, with correlation computed either with $r_\beta$ (*left*) and $R_\beta$ (*right*). (C) Sparsity of all neuronal subtypes in all 47 valid baseline networks and after introduction of additive noise of different levels. In (B–C), red lines indicate the respective median values.

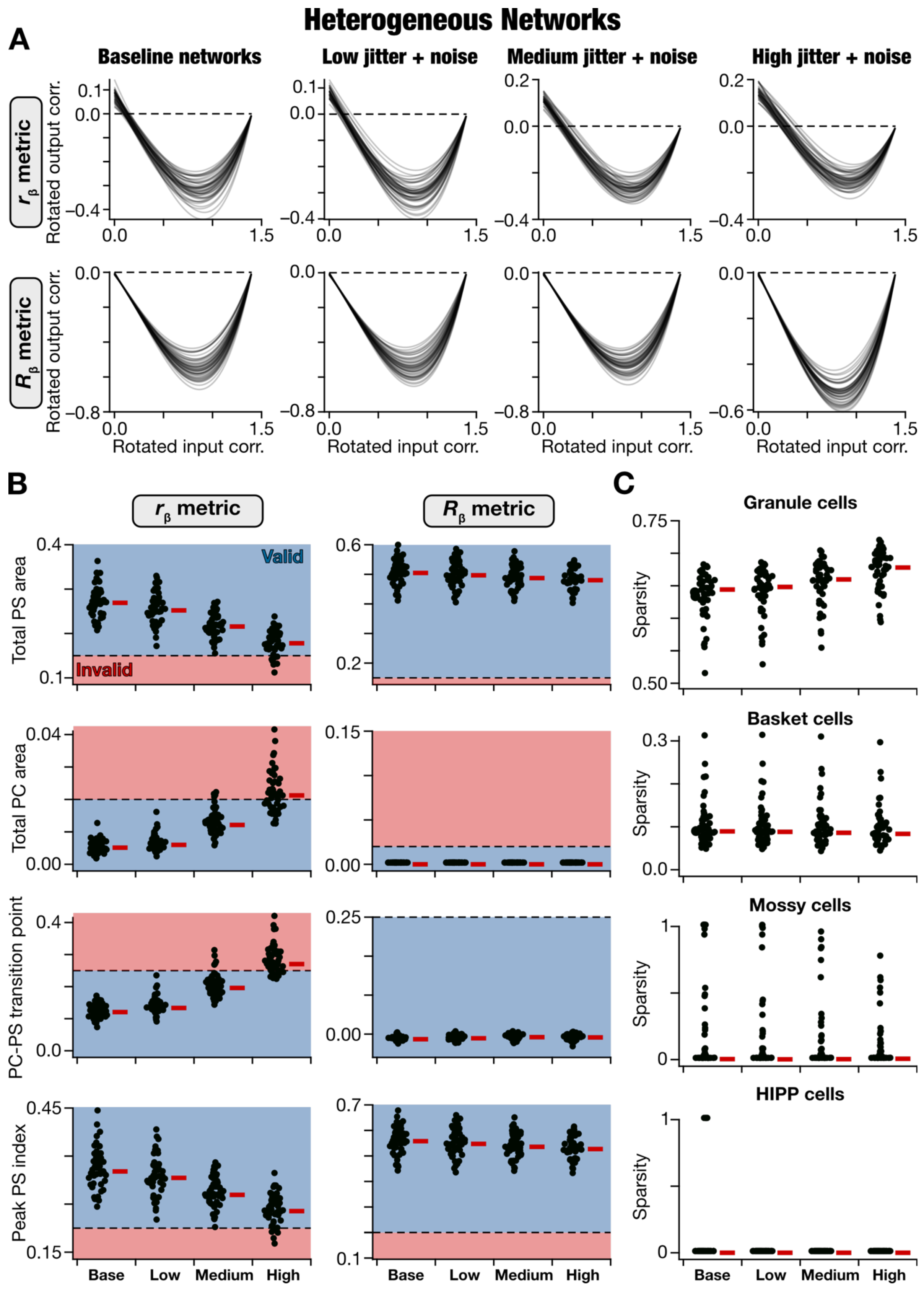

**Supplementary Figure S9: Network-to-network variability in the impact of introducing jitter and noise on pattern separation performance of the 47 valid heterogeneous DG networks.** (A) Visualization of pattern separation performance in all 47 valid baseline heterogeneous networks, and in each of the 47 networks with low, medium, and high levels of jitter and noise. Shown are fitted cubic polynomial on the rotated output correlation *vs*. rotated input correlation datapoints for all 11 morphed patterns. Correlation was computed either with the $r_\beta$ metric (*top*) or the $R_\beta$ metric (*bottom*). The pronounced variability in the impact of jitter and noise might be observed across networks. (B) All 4 pattern separation measurements in the valid networks and in networks where different levels of jitter and noise were introduced. These measurements were calculated from the traces shown in panel A for all 47 networks, with correlation computed either with $r_\beta$ (*left*) and $R_\beta$ (*right*). (C) Sparsity of all neuronal subtypes in all 47 valid baseline networks and after introduction of jitter and noise of different levels. In (B–C), red lines indicate the respective median values.

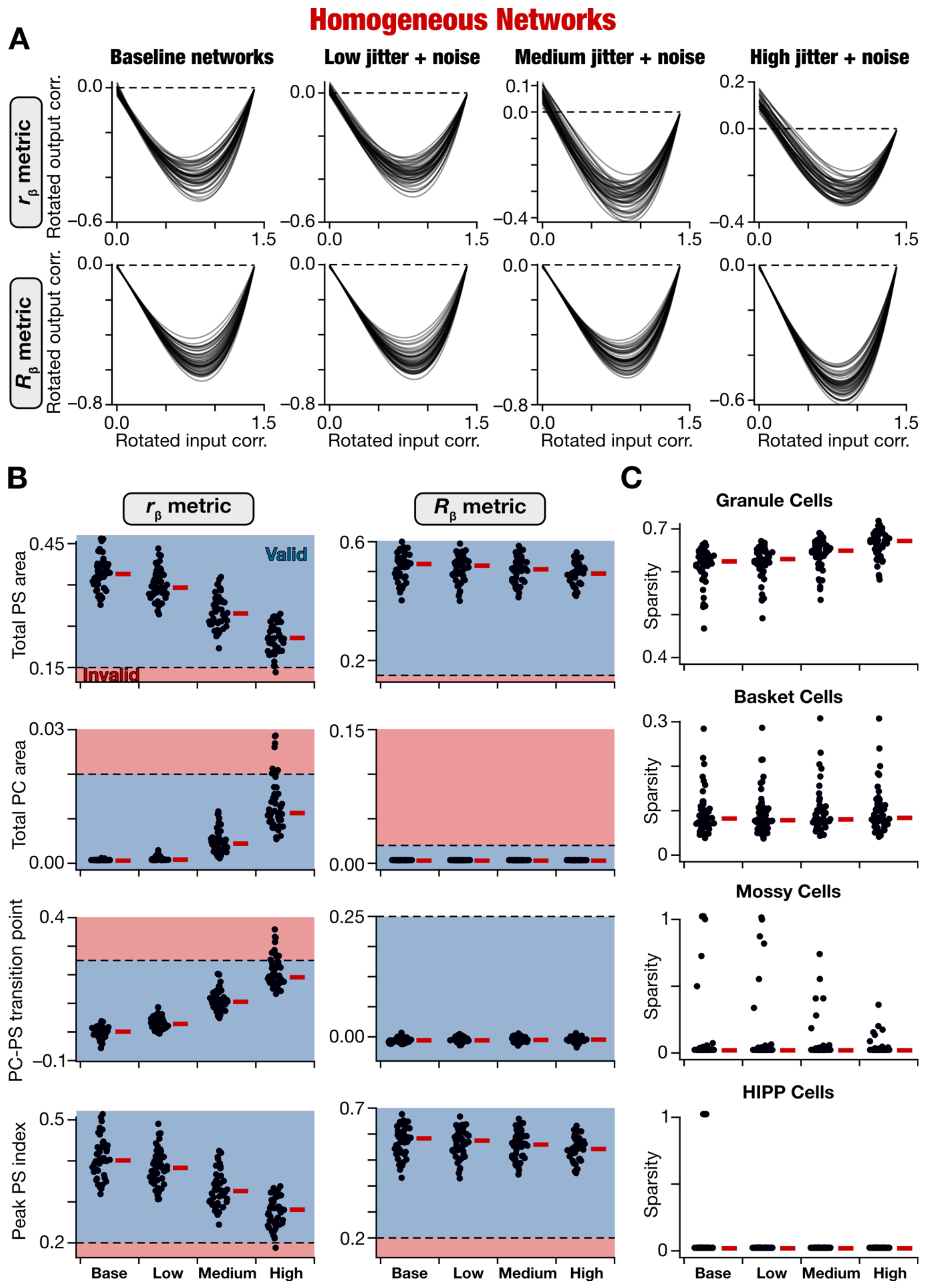

**Supplementary Figure S10: Network-to-network variability in the impact of introducing jitter and noise on pattern separation performance of the 47 valid homogeneous DG networks.** (A) Visualization of pattern separation performance in all 47 valid baseline homogeneous networks, and in each of the 47 networks with low, medium, and high levels of jitter and noise. Shown are fitted cubic polynomial on the rotated output correlation *vs*. rotated input correlation datapoints for all 11 morphed patterns. Correlation was computed either with the $r_\beta$ metric (*top*) or the $R_\beta$ metric (*bottom*). The pronounced variability in the impact of jitter and noise might be observed across networks. (B) All 4 pattern separation measurements in the valid networks and in networks where different levels of jitter and noise were introduced. These measurements were calculated from the traces shown in panel A for all 47 networks, with correlation computed either with $r_\beta$ (*left*) and $R_\beta$ (*right*). (C) Sparsity of all neuronal subtypes in all 47 valid baseline networks and after introduction of jitter and noise of different levels. In (B–C), red lines indicate the respective median values.

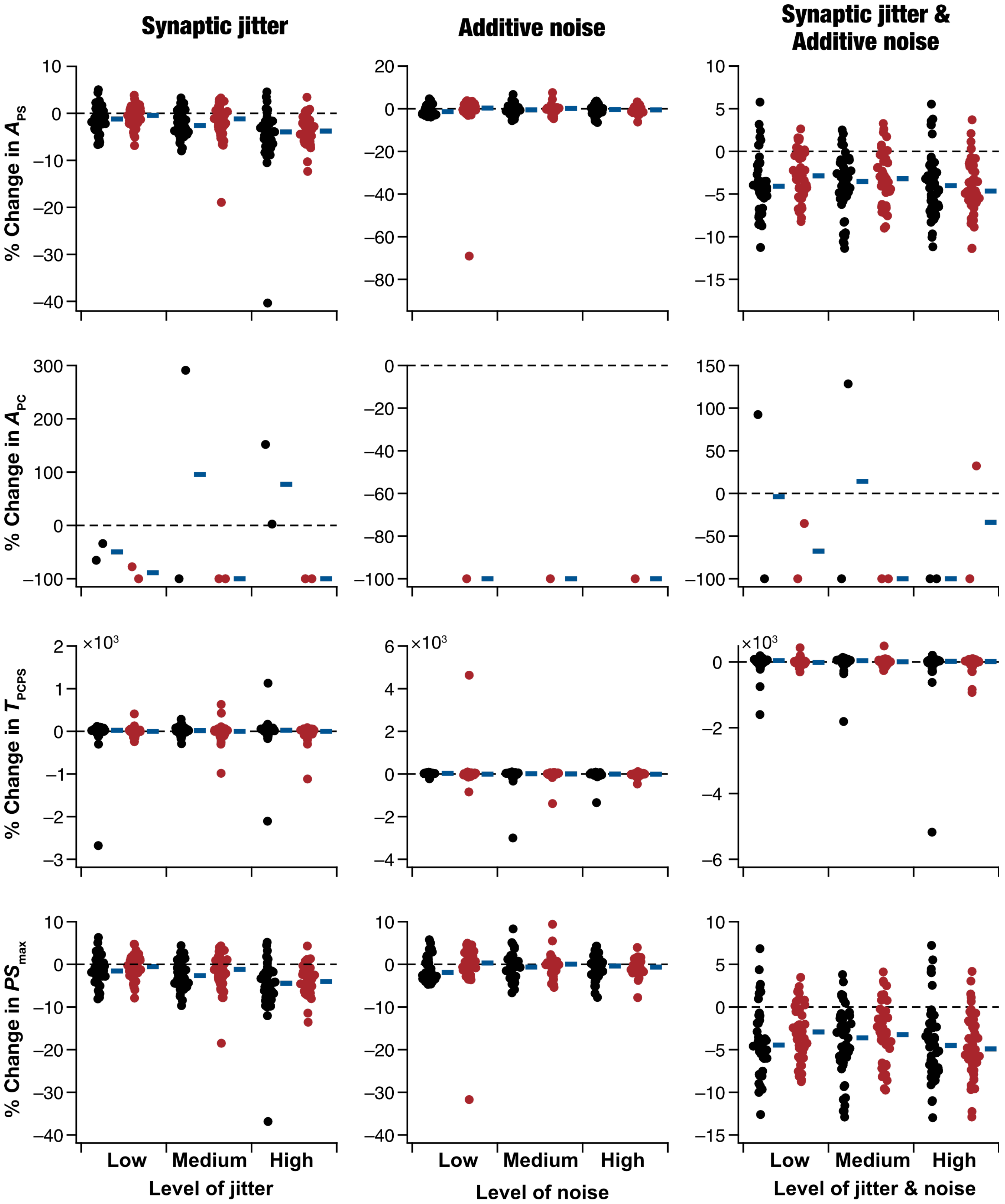

**Supplementary Figure S11: Percentage changes in pattern separation measurements computed using $R_\beta$ in heterogeneous and the homogeneous network challenged with perturbations.** Percentage changes in pattern separation measurements (computed using the $R_\beta$ metric) when perturbations of different levels were introduced. Percentage changes were computed with reference to the base networks where there were no perturbations. Perturbations were introduced in three distinct levels (low, medium, and high) as synaptic jitter (*left column*), or as additive noise to synaptic currents (*middle column*) or a combination of both jitter and noise (*right column*). The pronounced network-to-network variability, in both homogeneous and heterogeneous networks, in percentage changes of pattern separation measurements for the same levels of jitter and/or noise may be noted. In all plots, blue lines indicate the respective median values. The changes in the different measurements for the heterogeneous *vs.* homogeneous networks were not significantly different (Wilcox rank sum test).

**Supplementary Table S1. Base values and ranges for all the parameters used for generating a heterogeneous population for granule cells.**

| Granule cells | | | | |
|---|---|---|---|---|
| **Symbol** | **Parameter** | **Base Model** | **Lower Bound** | **Upper Bound** |
| $\boldsymbol{E_l\ (mV)}$ | Resting Potential | –75 | –77 | –73 |
| $\boldsymbol{R_m\ (k\Omega\ cm^2)}$ | Specific membrane resistance | 38 | 30 | 42 |
| $\boldsymbol{C_m\ (\mu F/cm^2\ )}$ | Specific membrane capacitance | 1 | 1 | 1 |
| $\boldsymbol{\tau_w\ (ms)}$ | Adaptation time constant | 300 | 200 | 500 |
| $\boldsymbol{\alpha\ (nS)}$ | Adaptation coupling parameter | 0.40 | 0 | 3 |
| $\boldsymbol{b\ (pA)}$ | Spike trigger parameter | 15 | 5 | 25 |
| $\boldsymbol{\Delta_T\ (mV)}$ | Slope Factor | 5 | 0 | 10 |
| $\boldsymbol{V_{Reset}\ (mV)}$ | Reset Voltage | –77 | –80 | –75 |
| $\boldsymbol{V_{th}\ (mV)}$ | Threshold voltage | –60 | –65 | –57 |
| $\boldsymbol{A\ (\mu m^2)}$ | Area | 30 | 20 | 40 |

**Supplementary Table S2. Base values and ranges for all the parameters used for generating a heterogeneous population for basket cells.**

| Basket cells | | | | |
|---|---|---|---|---|
| **Symbol** | **Parameter** | **Base Model** | **Lower Bound** | **Upper Bound** |
| $\boldsymbol{E_l\ (mV)}$ | Resting Potential | –65 | –67 | –63 |
| $\boldsymbol{R_m\ (k\Omega\ cm^2)}$ | Specific membrane resistance | 8 | 5 | 10 |
| $\boldsymbol{C_m\ (\mu F/cm^2\ )}$ | Specific membrane capacitance | 1 | 1 | 1 |
| $\boldsymbol{\tau_w\ (ms)}$ | Adaptation time constant | 80 | 20 | 100 |
| $\boldsymbol{\alpha\ (nS)}$ | Adaptation coupling parameter | 1.5 | 0 | 3 |
| $\boldsymbol{b\ (pA)}$ | Spike trigger parameter | 2 | 0 | 8 |
| $\boldsymbol{\Delta_T\ (mV)}$ | Slope Factor | 2 | 0 | 4 |
| $\boldsymbol{V_{Reset}\ (mV)}$ | Reset Voltage | –70 | –73 | –65 |
| $\boldsymbol{V_{th}\ (mV)}$ | Threshold voltage | –57 | –63 | –57 |
| $\boldsymbol{A\ (\mu m^2)}$ | Area | 13 | 10 | 14 |

**Supplementary Table S3. Base values and ranges for all the parameters used for generating a heterogeneous population for mossy cells.**

| Mossy cells | | | | |
|---|---|---|---|---|
| **Symbol** | **Parameter** | **Base Model** | **Lower Bound** | **Upper Bound** |
| $E_l\ (mV)$ | Resting Potential | –61 | –63 | –58 |
| $R_m\ (k\Omega\ cm^2)$ | Specific membrane resistance | 43 | 41 | 48 |
| $C_m\ (\mu F/cm^2)$ | Specific membrane capacitance | 1 | 1 | 1 |
| $\tau_w\ (ms)$ | Adaptation time constant | 400 | 100 | 500 |
| $\alpha\ (nS)$ | Adaptation coupling parameter | 0.8 | 0 | 3 |
| $b\ (pA)$ | Spike trigger parameter | 50 | 10 | 70 |
| $\Delta_T\ (mV)$ | Slope Factor | 1 | 0 | 10 |
| $V_{Reset}\ (mV)$ | Reset Voltage | –65 | –67 | –63 |
| $V_{th}\ (mV)$ | Threshold voltage | –47 | –40 | –32 |
| $A\ (\mu m^2)$ | Area | 17 | 10 | 18 |

**Supplementary Table S4. Base values and ranges for all the parameters used for generating a heterogeneous population for HIPP cells.**

| HIPP Cells | | | | |
|---|---|---|---|---|
| **Symbol** | **Parameter** | **Base Model** | **Lower Bound** | **Upper Bound** |
| $E_l\ (mV)$ | Resting Potential | –60 | –63 | –58 |
| $R_m\ (k\Omega\ cm^2)$ | Specific membrane resistance | 24 | 22 | 26 |
| $C_m\ (\mu F/cm^2)$ | Specific membrane capacitance | 1 | 1 | 1 |
| $\tau_w\ (ms)$ | Adaptation time constant | 115 | 50 | 300 |
| $\alpha\ (nS)$ | Adaptation coupling parameter | 0.6 | 0 | 3 |
| $b\ (pA)$ | Spike trigger parameter | 40 | 10 | 70 |
| $\Delta_T\ (mV)$ | Slope Factor | 2 | 0 | 10 |
| $V_{Reset}\ (mV)$ | Reset Voltage | –55 | –58 | –53 |
| $V_{th}\ (mV)$ | Threshold voltage | –42 | –45 | –39 |
| $A\ (\mu m^2)$ | Area | 4 | 3 | 10 |

**Supplementary Table S5. Bounds for the physiologically relevant measurements for all four neuronal subtypes.**

| Symbol | Measurement | Lower Bound | Upper Bound |
|---|---|---|---|
| **Granule Cells** | | | |
| $\tau_m$ $(ms)$ | Membrane time constant | 26 | 35 |
| $Sag$ | Sag ratio | 0.9 | 1 |
| $R_{in}$ $(M\Omega)$ | Input resistance | 107 | 228 |
| $f_{50}$ $(Hz)$ | Firing frequency when 50 pA current is injected | 0 | 0 |
| $f_{150}$ $(Hz)$ | Firing frequency when 150 pA current is injected | 10 | 15 |
| $SFA$ | Spike frequency adaptation | 0.1 | 0.8 |
| **Basket Cells** | | | |
| $\tau_m$ $(ms)$ | Membrane time constant | 6 | 10 |
| $Sag$ | Sag ratio | 0.9 | 1 |
| $R_{in}$ $(M\Omega)$ | Input resistance | 45 | 65 |
| $f_{50}$ $(Hz)$ | Firing frequency when 50 pA current is injected | 0 | 0 |
| $f_{150}$ $(Hz)$ | Firing frequency when 150 pA current is injected | 30 | 50 |
| $SFA$ | Spike frequency adaptation | 0.85 | 1 |
| **Mossy Cells** | | | |
| $\tau_m$ $(ms)$ | Membrane time constant | 26 | **32** |
| $Sag$ | Sag ratio | 0.6 | 0.91 |
| $R_{in}$ $(M\Omega)$ | Input resistance | 180 | 250 |
| $f_{50}$ $(Hz)$ | Firing frequency when 50 pA current is injected | 0 | 0 |
| $f_{150}$ $(Hz)$ | Firing frequency when 150 pA current is injected | 5 | 10 |
| $SFA$ | Spike frequency adaptation | 0.2 | 0.8 |
| **HIPP Cells** | | | |
| $\tau_m$ $(ms)$ | Membrane time constant | 14 | 20 |
| $Sag$ | Sag ratio | 0.5 | 0.95 |
| $R_{in}$ $(M\Omega)$ | Input resistance | 320 | 420 |
| $f_{50}$ $(Hz)$ | Firing frequency when 50 pA current is injected | 0 | 5 |
| $f_{150}$ $(Hz)$ | Firing frequency when 150 pA current is injected | 20 | 30 |
| $SFA$ | Spike frequency adaptation | 0.05 | 0.8 |

**Supplementary Table S6: Connection probability of different synaptic connections in the network. Rows represent presynaptic and columns represent postsynaptic cell (Chavlis et al., 2017).**

| | | Postsynaptic cells | | | | |
|---|---|---|---|---|---|---|
| | | PP | Granule | HIPP | Mossy | Basket |
| Presynaptic cells | PP | – | 0.015 | 0.035 | – | – |
| | Granule | – | – | – | 0.02 | 0.018 |
| | HIPP | – | 0.12 | – | – | – |
| | Mossy | – | 0.05 | – | – | 0.05 |
| | Basket | – | 0.16 | – | – | – |

**Supplementary Table S7: Rise and decay time constant along with synaptic reversal potential for all the synapses in the network, synaptic weights and axon delays (Geiger et al., 1997; Bartos et al., 2001; Larimer and Strowbridge, 2008; Myers and Scharfman, 2009; Chavlis et al., 2017).**

| Synapse (Pre → Post) | $\tau_{rise}$ (ms) | $\tau_{decay}$ (ms) | $E_{syn}$ (mV) | $\bar{g}_{syn}$ (nS) | Axon delay (ms) |
|---|---|---|---|---|---|
| **AMPA** | | | | | |
| PP → GC | 0.1 | 2.5 | 0 | 6.43 | 3.0 |
| GC → MC | 0.5 | 6.5 | 0 | 2.85 | 1.5 |
| GC → BC | 2.5 | 3.5 | 0 | 5.47 | 0.8 |
| PP → HIPP | 2.0 | 11.0 | 0 | 1.98 | 3.0 |
| MC → GC | 0.1 | 2.5 | 0 | 4.35 | 3.0 |
| MC → BC | 2.5 | 3.5 | 0 | 2.13 | 3.0 |
| **NMDA** | | | | | |
| PP → GC | 0.33 | 50.0 | 0 | 6.944 | 3.0 |
| PP → HIPP | 4.8 | 110.0 | 0 | 2.138 | 3.0 |
| GC → MC | 4.0 | 100.0 | 0 | 3.078 | 1.5 |
| MC → GC | 0.33 | 55.0 | 0 | 4.698 | 3.0 |
| MC → BC | 10.0 | 130.0 | 0 | 2.300 | 3.0 |
| GC → BC | 10.0 | 130.0 | 0 | 5.852 | 0.8 |
| **GABA** | | | | | |
| BC → GC | 0.9 | 6.8 | –90 | 4.29 | 0.85 |
| HIPP → GC | 0.9 | 6.8 | –90 | 2.65 | 1.6 |

**Supplementary Table S8: Parametric bounds for the 8 synaptic weights for running MPMOSS to obtain valid DG networks.**

| Pre → Post | Symbol | Lower Bound (nS) | Upper Bound (nS) |
|---|---|---|---|
| **PP → Granule** | $W_{\mathrm{GP}}$ | 0 | 20 |
| **PP → HIPP** | $W_{\mathrm{HP}}$ | 0 | 10 |
| **HIPP → Granule** | $W_{\mathrm{GH}}$ | 0 | 10 |
| **Granule → Mossy** | $W_{\mathrm{MG}}$ | 0 | 10 |
| **Granule → Basket** | $W_{\mathrm{BG}}$ | 0 | 10 |
| **Mossy → Granule** | $W_{\mathrm{GM}}$ | 0 | 10 |
| **Mossy → Basket** | $W_{BM}$ | 0 | 10 |
| **Basket → Granule** | $W_{\mathrm{GB}}$ | 0 | 10 |

**Supplementary Table S9: Measurements bounds used for the validation of networks that were effective in performing pattern separation.**

| Measurement | Description | Range |
|---|---|---|
| $\boldsymbol{A}_{\mathbf{PS}}$ | Total pattern separation area | $> 0.15$ |
| $\boldsymbol{A}_{\mathbf{PC}}$ | Total pattern completion area | $< 0.02$ |
| $\boldsymbol{T}_{\mathbf{PCPS}}$ | Pattern completion to pattern separation transition point | $< 0.25$ |
| $\boldsymbol{PS}_{\mathbf{max}}$ | Peak pattern separation index | $> 0.2$ |

**Supplementary Table S10: Outcomes of statistical tests for data in Figure 7.**

| Panel | Measurement | Groups | Test used | p value |
|---|---|---|---|---|
| **Figure 7B** | $A_{PS}\ (r_\beta)$ | Base, BC⁻, MC⁻, HC⁻ | Kruskal Wallis | $< 2.2 \times 10^{-16}$ |
| **Figure 7B** | $A_{PS}\ (r_\beta)$ | Base *vs*. BC⁻<br>Base *vs*. MC⁻<br>Base *vs*. HC⁻ | Wilcox signed rank | $1.4 \times 10^{-14}$<br>0.20<br>0.02 |
| **Figure 7B** | $A_{PC}\ (r_\beta)$ | Base, BC⁻, MC⁻, HC⁻ | Kruskal Wallis | $1.6 \times 10^{-13}$ |
| **Figure 7B** | $A_{PC}\ (r_\beta)$ | Base *vs*. BC⁻<br>Base *vs*. MC⁻<br>Base *vs*. HC⁻ | Wilcox signed rank | $1.4 \times 10^{-14}$<br>0.16<br>$8.2 \times 10^{-3}$ |
| **Figure 7B** | $T_{PCPS}\ (r_\beta)$ | Base, BC⁻, MC⁻, HC⁻ | Kruskal Wallis | $3.4 \times 10^{-8}$ |
| **Figure 7B** | $T_{PCPS}\ (r_\beta)$ | Base *vs*. BC⁻<br>Base *vs*. MC⁻<br>Base *vs*. HC⁻ | Wilcox signed rank | $1.5 \times 10^{-5}$<br>$5.3 \times 10^{-4}$<br>$1.8 \times 10^{-4}$ |
| **Figure 7B** | $PS_{\max}\ (r_\beta)$ | Base, BC⁻, MC⁻, HC⁻ | Kruskal Wallis | $< 2.2 \times 10^{-16}$ |
| **Figure 7B** | $PS_{\max}\ (r_\beta)$ | Base *vs*. BC⁻<br>Base *vs*. MC⁻<br>Base *vs*. HC⁻ | Wilcox signed rank | $1.4 \times 10^{-14}$<br>0.08<br>$6.7 \times 10^{-3}$ |
| **Figure 7B** | $A_{PS}\ (R_\beta)$ | Base, BC⁻, MC⁻, HC⁻ | Kruskal Wallis | $< 2.2 \times 10^{-16}$ |
| **Figure 7B** | $A_{PS}\ (R_\beta)$ | Base *vs*. BC⁻<br>Base *vs*. MC⁻<br>Base *vs*. HC⁻ | Wilcox signed rank | $2.8 \times 10^{-14}$<br>$6.2 \times 10^{-10}$<br>$3.0 \times 10^{-11}$ |
| **Figure 7B** | $A_{PC}\ (R_\beta)$ | Base, BC⁻, MC⁻, HC⁻ | Kruskal Wallis | $< 2.2 \times 10^{-16}$ |
| **Figure 7B** | $A_{PC}\ (R_\beta)$ | Base *vs*. BC⁻<br>Base *vs*. MC⁻<br>Base *vs*. HC⁻ | Wilcox signed rank | $5.7 \times 10^{-14}$<br>$6.5 \times 10^{-5}$<br>$4.5 \times 10^{-6}$ |
| **Figure 7B** | $T_{PCPS}\ (R_\beta)$ | Base, BC⁻, MC⁻, HC⁻ | Kruskal Wallis | $< 2.2 \times 10^{-16}$ |
| **Figure 7B** | $T_{PCPS}\ (R_\beta)$ | Base *vs*. BC⁻<br>Base *vs*. MC⁻<br>Base *vs*. HC⁻ | Wilcox signed rank | $5.8 \times 10^{-11}$<br>0.07<br>$5.4 \times 10^{-3}$ |
| **Figure 7B** | $PS_{\max}\ (R_\beta)$ | Base, BC⁻, MC⁻, HC⁻ | Kruskal Wallis | $< 2.2 \times 10^{-16}$ |
| **Figure 7B** | $PS_{\max}\ (R_\beta)$ | Base *vs*. BC⁻<br>Base *vs*. MC⁻<br>Base *vs*. HC⁻ | Wilcox signed rank | $5.4 \times 10^{-10}$<br>$1.8 \times 10^{-7}$<br>$2.9 \times 10^{-9}$ |
| **Figure 7C** | $Sparsity$ (GCs) | Base, BC⁻, MC⁻, HC⁻ | Kruskal Wallis | $< 2.2 \times 10^{-16}$ |
| **Figure 7C** | $Sparsity$ (GCs) | Base *vs*. BC⁻<br>Base *vs*. MC⁻<br>Base *vs*. HC⁻ | Wilcox signed rank | $2.5 \times 10^{-9}$<br>$7.1 \times 10^{-14}$<br>$7.1 \times 10^{-14}$ |
| **Figure 7C** | $Sparsity$ (BCs) | Base, MC⁻, HC⁻ | Kruskal Wallis | $< 2.2 \times 10^{-16}$ |
| **Figure 7C** | $Sparsity$ (BCs) | Base *vs*. MC⁻<br>Base *vs*. HC⁻ | Wilcox signed rank | $2.5 \times 10^{-9}$<br>$2.5 \times 10^{-9}$ |
| **Figure 7C** | $Sparsity$ (MCs) | Base, BC⁻, HC⁻ | Kruskal Wallis | $< 2.2 \times 10^{-16}$ |
| **Figure 7C** | $Sparsity$ (MCs) | Base *vs*. BC⁻<br>Base *vs*. HC⁻ | Wilcox signed rank | $1.2 \times 10^{-6}$<br>$1.2 \times 10^{-6}$ |
| **Figure 7C** | $Sparsity$ (HIPP) | Base, BC⁻, MC⁻ | Kruskal Wallis | 0.65 |
| **Figure 7C** | $Sparsity$ (HIPP) | Base *vs*. BC⁻<br>Base *vs*. MC⁻ | Wilcox signed rank | 0.18<br>0.18 |

**Supplementary Table S11: Outcomes of statistical tests for data in Figure 8.**

| Measurement | Groups | Test used | p value |
|---|---|---|---|
| **Synaptic jitter** | | | |
| **Change in $A_{PS}$** | Low jitter<br>Homogeneous *vs*. Heterogeneous | Wilcox rank sum | $1.0 \times 10^{-13}$ |
| **Change in $A_{PS}$** | Medium jitter<br>Homogeneous *vs*. Heterogeneous | Wilcox rank sum | $1.7 \times 10^{-16}$ |
| **Change in $A_{PS}$** | High jitter<br>Homogeneous *vs*. Heterogeneous | Wilcox rank sum | $7.3 \times 10^{-19}$ |
| **Change in $A_{PC}$** | Low jitter<br>Homogeneous *vs*. Heterogeneous | Wilcox rank sum | $3.3 \times 10^{-3}$ |
| **Change in $A_{PC}$** | Medium jitter<br>Homogeneous *vs*. Heterogeneous | Wilcox rank sum | $4.0 \times 10^{-16}$ |
| **Change in $A_{PC}$** | High jitter<br>Homogeneous *vs*. Heterogeneous | Wilcox rank sum | $4.0 \times 10^{-16}$ |
| **Change in $T_{PCPS}$** | Low jitter<br>Homogeneous *vs*. Heterogeneous | Wilcox rank sum | $7.0 \times 10^{-10}$ |
| **Change in $T_{PCPS}$** | Medium jitter<br>Homogeneous *vs*. Heterogeneous | Wilcox rank sum | $1.2 \times 10^{-27}$ |
| **Change in $T_{PCPS}$** | High jitter<br>Homogeneous *vs*. Heterogeneous | Wilcox rank sum | $1.2 \times 10^{-27}$ |
| **Change in $PS_{\max}$** | Low jitter<br>Homogeneous *vs*. Heterogeneous | Wilcox rank sum | $1.0 \times 10^{-6}$ |
| **Change in $PS_{\max}$** | Medium jitter<br>Homogeneous *vs*. Heterogeneous | Wilcox rank sum | $8.4 \times 10^{-10}$ |
| **Change in $PS_{\max}$** | High jitter<br>Homogeneous *vs*. Heterogeneous | Wilcox rank sum | $1.8 \times 10^{-12}$ |

**Supplementary Table S11: Outcomes of statistical tests for data in Figure 8 (contd).**

| Measurement | Groups | Test used | p value |
|---|---|---|---|
| **Additive noise** | | | |
| **Change in $A_{PS}$** | Low noise<br>Homogeneous *vs*. Heterogeneous | Wilcox rank sum | 0.28 |
| **Change in $A_{PS}$** | Medium noise<br>Homogeneous *vs*. Heterogeneous | Wilcox rank sum | 0.96 |
| **Change in $A_{PS}$** | High noise<br>Homogeneous *vs*. Heterogeneous | Wilcox rank sum | 0.94 |
| **Change in $A_{PC}$** | Low noise<br>Homogeneous *vs*. Heterogeneous | Wilcox rank sum | $2.9 \times 10^{-3}$ |
| **Change in $A_{PC}$** | Medium noise<br>Homogeneous *vs*. Heterogeneous | Wilcox rank sum | $< 2.2 \times 10^{-5}$ |
| **Change in $A_{PC}$** | High noise<br>Homogeneous *vs*. Heterogeneous | Wilcox rank sum | $1.2 \times 10^{-6}$ |
| **Change in $T_{PCPS}$** | Low noise<br>Homogeneous *vs*. Heterogeneous | Wilcox rank sum | 0.76 |
| **Change in $T_{PCPS}$** | Medium noise<br>Homogeneous *vs*. Heterogeneous | Wilcox rank sum | 0.43 |
| **Change in $T_{PCPS}$** | High noise<br>Homogeneous *vs*. Heterogeneous | Wilcox rank sum | 0.91 |
| **Change in $PS_{\max}$** | Low noise<br>Homogeneous *vs*. Heterogeneous | Wilcox rank sum | 0.20 |
| **Change in $PS_{\max}$** | Medium noise<br>Homogeneous *vs*. Heterogeneous | Wilcox rank sum | 0.98 |
| **Change in $PS_{\max}$** | High noise<br>Homogeneous *vs*. Heterogeneous | Wilcox rank sum | 0.43 |

**Supplementary Table S11: Outcomes of statistical tests for data in Figure 8 (contd).**

| Measurement | Groups | Test used | p value |
|---|---|---|---|
| **Synaptic jitter & Additive noise** | | | |
| **Change in $A_{PS}$** | Low jitter & noise<br>Homogeneous *vs*. Heterogeneous | Wilcox rank sum | 0.03 |
| **Change in $A_{PS}$** | Medium jitter & noise<br>Homogeneous *vs*. Heterogeneous | Wilcox rank sum | $4.1 \times 10^{-5}$ |
| **Change in $A_{PS}$** | High jitter & noise<br>Homogeneous *vs*. Heterogeneous | Wilcox rank sum | $1.6 \times 10^{-5}$ |
| **Change in $A_{PC}$** | Low jitter & noise<br>Homogeneous *vs*. Heterogeneous | Wilcox rank sum | 0.04 |
| **Change in $A_{PC}$** | Medium jitter & noise<br>Homogeneous *vs*. Heterogeneous | Wilcox rank sum | $8.4 \times 10^{-13}$ |
| **Change in $A_{PC}$** | High jitter & noise<br>Homogeneous *vs*. Heterogeneous | Wilcox rank sum | $4.0 \times 10^{-16}$ |
| **Change in $T_{PCPS}$** | Low jitter & noise<br>Homogeneous *vs*. Heterogeneous | Wilcox rank sum | 0.04 |
| **Change in $T_{PCPS}$** | Medium jitter & noise<br>Homogeneous *vs*. Heterogeneous | Wilcox rank sum | $7.1 \times 10^{-16}$ |
| **Change in $T_{PCPS}$** | High jitter & noise<br>Homogeneous *vs*. Heterogeneous | Wilcox rank sum | $9.0 \times 10^{-24}$ |
| **Change in $PS_{\max}$** | Low jitter & noise<br>Homogeneous *vs*. Heterogeneous | Wilcox rank sum | 0.09 |
| **Change in $PS_{\max}$** | Medium jitter & noise<br>Homogeneous *vs*. Heterogeneous | Wilcox rank sum | $2.9 \times 10^{-4}$ |
| **Change in $PS_{\max}$** | High jitter & noise<br>Homogeneous *vs*. Heterogeneous | Wilcox rank sum | $7.7 \times 10^{-5}$ |
